# The metabolic cofactor Coenzyme A enhances alternative macrophage activation via MyD88-linked signaling

**DOI:** 10.1101/2024.03.28.587096

**Authors:** Anthony E. Jones, Amy Rios, Neira Ibrahimovic, Carolina Chavez, Nicholas A. Bayley, Andréa B. Ball, Wei Yuan Hsieh, Alessandro Sammarco, Amber R. Bianchi, Angel A. Cortez, Thomas G. Graeber, Alexander Hoffmann, Steven J. Bensinger, Ajit S. Divakaruni

**Affiliations:** Departments of Molecular and Medical Pharmacology, University of California, Los Angeles, Los Angeles, CA 90095, USA; Microbiology, Immunology, and Molecular Genetics, University of California, Los Angeles, Los Angeles, CA 90095, USA

## Abstract

Metabolites and metabolic co-factors can shape the innate immune response, though the pathways by which these molecules adjust inflammation remain incompletely understood. Here we show that the metabolic cofactor Coenzyme A (CoA) enhances IL-4 driven alternative macrophage activation [m(IL-4)] *in vitro* and *in vivo*. Unexpectedly, we found that perturbations in intracellular CoA metabolism did not influence m(IL-4) differentiation. Rather, we discovered that exogenous CoA provides a weak TLR4 signal which primes macrophages for increased receptivity to IL-4 signals and resolution of inflammation via MyD88. Mechanistic studies revealed MyD88-linked signals prime for IL-4 responsiveness, in part, by reshaping chromatin accessibility to enhance transcription of IL-4-linked genes. The results identify CoA as a host metabolic co-factor that influences macrophage function through an extrinsic TLR4-dependent mechanism, and suggests that damage-associated molecular patterns (DAMPs) can prime macrophages for alternative activation and resolution of inflammation.

## INTRODUCTION

Macrophages are innate immune cells that execute a variety of functions such as detecting and removing foreign pathogens, instructing adaptive immune cell function, secreting cytokines, and maintaining tissue homeostasis. To fulfill these diverse roles, macrophages link the detection of distinct external cues with the engagement of specific transcriptional programs to support different functional states. This process is commonly termed macrophage activation or ‘polarization’^1,2^.

*In vitro* studies often consider macrophage polarization in two discrete states: classical (Type 1) and alternative (Type 2) activation^3^. Classical activation is associated with antimicrobial immunity, and occurs when macrophages detect pathogen-associated molecular patterns (PAMPs) such as lipopolysaccharide (LPS) or damage-associated molecular patterns (DAMPs). Alternative activation is associated with wound healing along with allergen and helminth immunity, and is initiated by the macrophage response to interleukin-4 (IL-4) ± IL-13.

Along with transcriptional programs that support core macrophage functions such as cytokine secretion and phagocytosis, changes in cellular metabolism are also important and perhaps essential for macrophage effector function^4–6^. Upon classical activation with LPS, macrophages increase glycolysis and repurpose mitochondria away from oxidative phosphorylation and towards the generation of metabolic signals thought to amplify the pro-inflammatory response^7–10^. Conversely, alternatively activated macrophages (AAMs) display increased rates of oxidative phosphorylation and fatty acid oxidation^11,12^, though evidence is mixed as to whether these changes are essential for the macrophage IL-4 response [m(IL-4)]^13–15^. The mechanistic links between how increased oxidative phosphorylation and/or fatty acid oxidation could specifically support m(IL-4) are also unclear. Proposed mechanisms, though, include changes in histone acetylation from enhanced acetyl CoA production^16^ as well as transcriptional changes that respond to the mitochondrial membrane potential^17^

Previous work in our laboratory has associated intracellular levels of CoA with the macrophage IL-4 response^13,18^. When trying to identify why excess concentrations of the CPT-1 inhibitor etomoxir blocked m(IL-4) but genetic ablation of either *Cpt1* or *Cpt2* did not^14,19^, we discovered that excess etomoxir disrupted macrophage CoA homeostasis. In support of this as a putative mechanism, provision of exogenous CoA restored both intracellular CoA levels as well as the expression of AAM-associated cell surface markers^13^. However, precisely how CoA instructs macrophage activation was not studied.

Here we demonstrate that CoA augments AAM function via weak toll-like receptor 4 (TLR4) agonism and myeloid differentiation primary response protein 88 (MyD88)-linked signaling, a pathway commonly associated with classical or pro-inflammatory activation. In investigating the mechanism by which exogenous CoA regulates m(IL-4), we surprisingly discovered that CoA provision did not act either by changing intracellular CoA levels or by enhancing known metabolic hallmarks of the IL-4 response. Rather, pharmacologic and genetic approaches showed exogenous CoA is a weak TLR4 agonist and boosts m(IL-4) by activating MyD88-linked signaling. MyD88 agonism was sufficient to enhance *in vitro* and *in vivo* alternative activation and increase chromatin accessibility at the promoter regions of IL-4-target genes. The data show that (i) CoA is a TLR4 agonist, (ii) many of the metabolic hallmarks of the m(IL-4) do not always correlate with anti-inflammatory activation, and (iii) pro-inflammatory MyD88-linked signaling can support AAM function and resolution of inflammation. Furthermore, the results indicate CoA can act as a damage-associated molecular pattern (DAMP) that primes macrophages for the resolution of inflammation by an extrinsic TLR4-dependent mechanism.

## RESULTS

### Exogenous CoA provision enhances alternative macrophage activation in vitro and in vivo

Prior work had shown that exogenous CoA could rescue the inhibition of m(IL-4) by etomoxir, but the mechanisms underlying this effect were not defined. To better understand how exogenous CoA influenced alternative macrophage activation, mouse bone marrow-derived macrophages (BMDMs) were treated with IL-4 alone or in combination with CoA for 48 hr. Gene expression studies revealed that CoA enhanced the expression of multiple IL-4-associated genes, including *Mgl2, Pdcd1gl2, Fizz1, Chil4, Ccl8, Arg1, and Mrc1* (Figure 1a and Supplemental Fig 1a)^17,20–22^. Flow cytometry analysis revealed a similar relationship when measuring IL-4-linked cell surface marker expression. CoA increased the mean fluorescence intensity of CD206 and CD301, as well as the frequency of CD206^+^/CD71^+^ and CD206^+^/CD301^+^ BMDMs (Figs. 1b-d). This effect was observed with CoA concentrations as low as 62.5µM (Supplemental Fig. S1b). Importantly, CoA itself did not stimulate the expression of IL-4-associated genes or cell surface markers (Figs. 1a-d), demonstrating it is not an IL-4 receptor agonist but rather acts cooperatively to enhance AAM differentiation.

**Figure 1.**
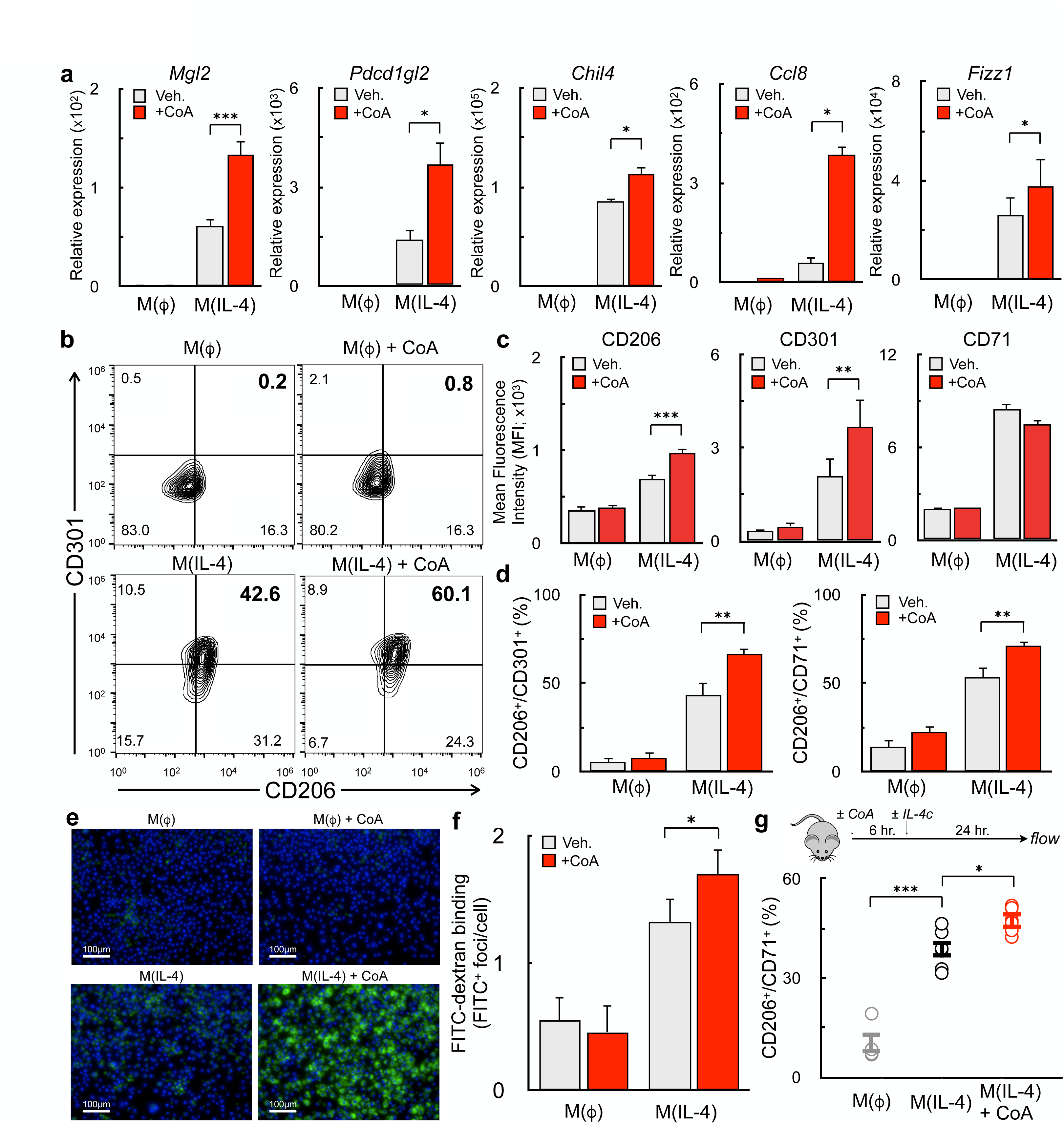
Exogenous CoA provision enhances alternative macrophage activation. **(a)** qPCR analysis of the IL-4-associated genes *Mgl2*, *Pdcd1gl2*, *Chil4*, *Ccl8,* and *Fizz1* in BMDMs treated with CoA (1 mM), IL-4 (20 ng/mL), CoA + IL-4, or vehicle for 48 hr. (n≥9 independent biological replicates). **(b-d)** Flow cytometric analysis of the IL-4-associated cell surface markers CD206, CD301, and CD71 after treatments as in (a). (b) Contour plots with the percentage of cells expressing both CD206 and CD301 is indicated in the upper right quadrant. Data shown are from a single representative experiment. (c) Aggregate mean fluorescence intensity of CD206, CD301, and CD71 (n=9 independent biological replicates). (d) Percentage of CD206^+^/CD301^+^ and CD206^+^/CD71^+^ populations (n=9 independent biological replicates). **(e)** Representative images of BMDMs incubated for 1 hr. with FITC-Dextran (1mg/mL, green) and Hoechst 3342 (10ng/mL, blue) after stimulation with compounds as in (a). **(f)** Aggregate image analysis data for experiments as in (e) (n=3 independent biological replicates). **(G)** Quantification of CD206^+^/CD71^+^ peritoneal macrophages from mice that were exposed to vehicle (PBS), CoA (40mg/kg), IL-4c (5 µg IL-4 and 25 µg anti-IL-4 monoclonal antibody), or the combination of IL-4c + CoA. (n≥3 mice were used for each group). All data are presented as mean ± SEM. *p < 0.05; **p < 0.01; ***p < 0.001.

Mannose receptor activity is essential for the initiation of the T_H_2 response during the helminth infection^23,24^. Given the observed increase in its gene (*Mrc1*) and protein (CD206) expression, we measured the effect of exogenous CoA on activity of the mannose receptor. As expected, high-content imaging revealed CoA enhanced the cellular uptake of FITC-dextran, a fluorescently labeled polysaccharide and mannose receptor ligand^25^ (Figs. 1e&f).

To assess whether CoA could enhance alternative activation *in vivo,* mice were injected with IL-4 complex (IL-4c) i.p. in the presence or absence of CoA (40 mg/kg). After one day, the peritoneum was flushed and the frequency of CD206^+^/CD71^+^ peritoneal macrophages assessed. Indeed, CoA enhanced the fraction of AAMs co-expressing CD206 and CD71 (Figure 1g), demonstrating that it enhances IL-4-mediated AAM differentiation *in vitro* and *in vivo*.

### CoA does not augment alternative macrophage activation by enhancing metabolic hallmarks of M(IL-4)

We next sought to identify the mechanism by which CoA enhances m(IL-4). Several metabolic hallmarks of AAMs require CoA as a necessary cofactor, including enhanced mitochondrial respiratory capacity^26–28^, mitochondrial pyruvate oxidation^12,29^, and *de novo* lipid synthesis^30,31^. As such, we hypothesized that addition of exogenous CoA enhanced the IL-4 response by increasing intracellular CoA levels to support flux through these metabolic pathways^32–34^.

We first confirmed that exogenously added CoA could expand the cellular CoA pool. Supplementing culture medium with CoA increased the steady-state abundance of both intracellular CoA and acetyl CoA (Fig. 2a). Next, we investigated whether CoA provision enhanced IL-4-driven increases in respiration and glycolysis induced by IL-4^12,26,30^. Similar to other reports, we observed increases in ATP-linked respiration, maximal respiratory capacity, and glycolysis with IL-4. However, addition of CoA did not further augment these metabolic changes, and even limited the maximal respiratory capacity of AAMs (Figs. 2b-d).

**Figure 2:**
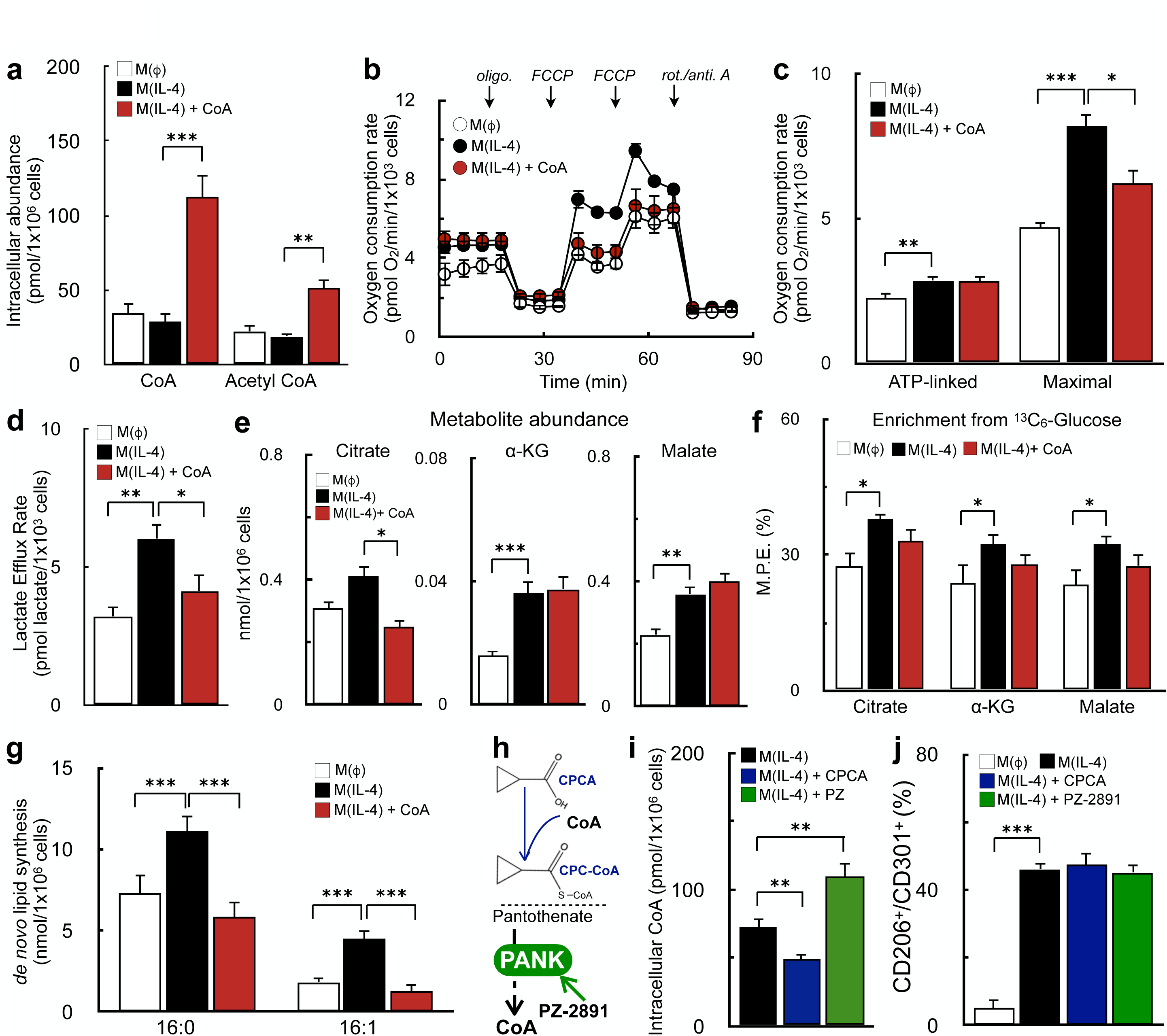
CoA does not enhance alternative macrophage activation by boosting known metabolic hallmarks of the IL-4 response. **(a)** Intracellular levels of CoA and acetyl CoA in BMDMs measured by LC-MS/MS. Cells were treated with IL-4 (20 ng/mL), IL-4 + CoA (1 mM), or vehicle for 48 hr. as in Fig. 1 (n=3 independent biological replicates). **(b)** Representative respirometry trace of BMDMs treated as in (a). (n=5 technical replicates from a single biological replicate). **(c)** Aggregate ATP-linked and FCCP-stimulated respiration in intact BMDMs for treatments as in (a). Cells were offered 8 mM glucose, 2 mM pyruvate, and 2 mM glutamine in the experimental medium (n=8 independent biological replicates). **(d)** Lactate efflux rate from respirometry experiments in (b & c) calculated using Seahorse XF data and correcting for respiratory CO_2_ (n=8 independent biological replicates). **(e)** Metabolite abundances of citrate, α-ketoglutarate(α-KG), and malate in BMDMs treated as in (a) (n=7 independent biological replicates). **(f)** Enrichment from uniformly labeled ^13^C_6_-glucose into the TCA cycle intermediates as in (e) (n=8 independent biological replicates). **(g)** Quantification of newly synthesized palmitic acid (16:0) and palmitoleic acid (16:1) from BMDMs stimulated as in (a). (data shown as n=8 technical replicates from n=2 independent biological replicates). **(h)** Schematic depicting the mechanism of action of cyclopropane carboxylic acid (CPCA) and PZ-2891**. (i)** Intracellular CoA levels of BMDMs stimulated with IL-4, IL-4 + CPCA (1 mM), or IL-4+PZ-2891 (10 µM) for 48 h (n=5 independent biological replicates). **(j)** Flow cytometric quantification of the CD206^+^/CD301^+^ population for BMDMs treated as in (i) (n=5 independent biological replicates). All data are presented as mean ± SEM. *p < 0.05; **p < 0.01; ***p < 0.001.

After determining that CoA does not enhance alternative macrophage activation via an expansion of bioenergetic capacity, we then examined whether CoA affected other IL-4-linked metabolic alterations such as increased abundance of TCA cycle metabolites^29^ or enhanced pyruvate oxidation^12^, glutamine oxidation^29^, and *de novo* lipogenesis^31^. Indeed, we reproduced previous reports that show IL-4 increases steady-state levels of select TCA cycle metabolites (Figs. 2e), enrichment from glucose into the TCA cycle (Fig. 2f), and *de novo* lipid synthesis (Fig. 2g). However, as before, addition of CoA did not further increase these metabolic changes (Figs. 2e-g, S2a-c), and even blocked IL-4-stimulated increases in lipogenesis (Figs. 2g, S2d). As such, the results show exogenous CoA augments alternative activation by a mechanism discrete from reprogramming metabolism.

### Alterations in intracellular CoA levels are not sufficient to alter M(IL-4)

We then questioned whether changes in intracellular CoA levels, in fact, shape the IL-4 response. To answer this, we utilized two compounds with opposing impacts on intracellular CoA levels. To decrease steady-state CoA levels, we treated IL-4-stimulated BMDMs with cyclopropane carboxylic acid (CPCA), which decreases the abundance of “free” CoA as its cognate thioester CPC-CoA is formed^18,35^ (Fig. 2h). To increase steady-state CoA levels, we treated IL-4-polarized BMDMs with PZ-2891, a pantothenate kinase agonist which relieves inhibition of CoA biosynthesis^36^ (Fig. 2h). As expected, CPCA decreased intracellular levels of CoA and acetyl CoA, while PZ-2891 increased their abundance (Fig. 2i, S2e). Surprisingly, despite altering steady-state intracellular CoA levels, neither compound impacted alternative activation (Fig. 2j). The results show that although exogenous CoA provision augments the macrophage IL-4 response, the mechanism cannot be attributed to changing intracellular levels of CoA and acetyl CoA.

### Exogenous CoA induces a macrophage pro-inflammatory response in vitro and in vivo

Neither metabolic alterations nor intracellular CoA levels could explain why CoA provision enhanced m(IL-4). Therefore, we sought a more complete understanding of how CoA impacts the transcriptome of AAMs by conducting bulk RNA sequencing (RNA-seq) on naïve macrophages alongside those that were stimulated with either IL-4 or IL-4 with CoA. As expected, cells stimulated only with IL-4 increased the expression of genes associated with alternative activation and decreased expression of genes associated with classical activation relative to vehicle controls. In line with qPCR studies (Fig. 1a), CoA provision further increased the expression of IL-4-linked genes associated with alternative activation (Fig. 3a, *right*). Unexpectedly, however, CoA addition also increased the expression of genes associated with classical activation that were not associated with the IL-4 response (Fig. 3a, *right*). Subsequent Gene Set Enrichment Analysis (GSEA) showed that genes associated with TLR signaling pathways were upregulated upon co-treatment with CoA and IL-4 (Fig. 3b).

**Figure 3.**
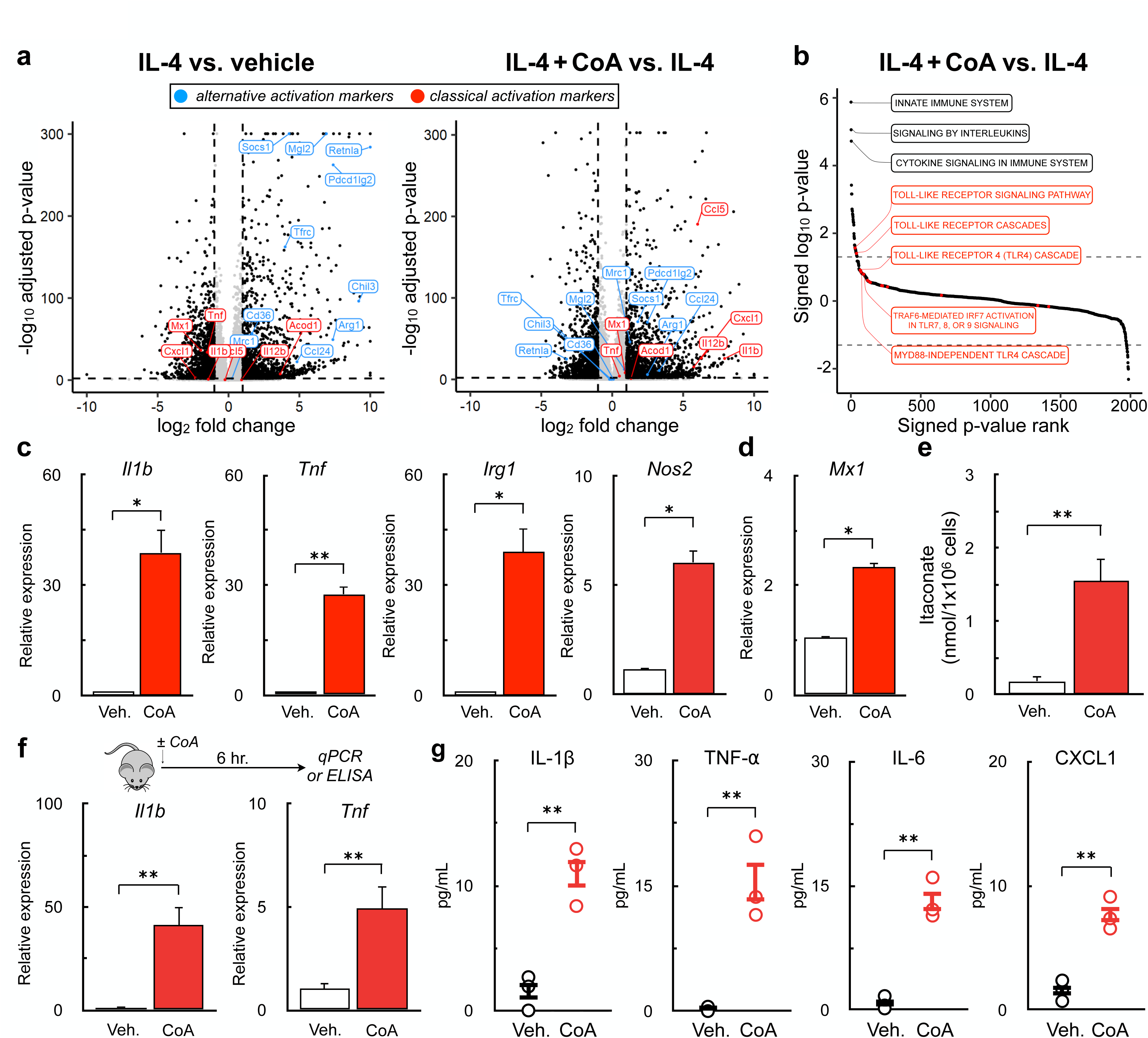
Exogenous CoA induces a pro-inflammatory response in BMDMs. **(a)** Volcano plot from bulk RNA sequencing data from BMDMs treated IL-4 (20 ng/mL), IL-4 + 1 mM CoA, or vehicle control for 48 hr. comparing differential gene expression between IL-4 vs. vehicle controls (left) and IL-4 + CoA vs. IL-4 (right). Genes associated with classical activation are depicted in red, genes associated with alternative activation are shown in blue. **(b)** Gene Set Enrichment Analysis of genes upregulated in BMDMs treated with IL-4+CoA vs. IL-4 alone. **(c)** qPCR analysis of *Il1b*, *Tnf*, *Irg1*, and *Nos2* in BMDMs stimulated with CoA (1mM) or vehicle control for 4 hr. (n=4 independent biological replicates). **(d)** qPCR analysis of the interferon-stimulated gene *Mx1* in BMDMs stimulated with 1 mM CoA or vehicle control for 24 hr. (n=4 independent biological replicates). **(e)** Itaconate abundance after treatment with 1 mM CoA or vehicle control for 48 hr. (n=6 independent biological replicates). **(f)** qPCR analysis of *Il1b* and *Tnf* in the peritoneal exudate cells of mice treated with (40 mg/kg) CoA 6 hr. prior to collection (n≥5 mice for each group). **(g)** Quantification of cytokines in the peritoneal lavage fluid (PLF) of mice treated as in (g) using Multiplexed ELISA (n=3 mice were used for each group). All data are presented as mean ± SEM. *p < 0.05; **p < 0.01.

This unbiased approach suggested that CoA may elicit a pro-inflammatory response in BMDMs along with its ability to enhance m(IL-4). To confirm this, we assessed whether CoA itself could induce expression of pro-inflammatory genes in the absence of IL-4. Indeed, exogenous CoA induced expression of *Il1b*, *Tnf*, *Nos2*, and *Irg1* (Fig. 3c), genes linked to the TLR adaptor protein MyD88, as well as the interferon-stimulated gene (ISG) *Mx1* (Fig. 3d). As *Irg1* encodes the enzyme generating the anti-microbial metabolite itaconate, we also observed a ∼10-fold increase in itaconate synthesis upon CoA provision (Fig. 3e). Lastly, i.p. administration of CoA in mice increased expression of *Il1b* and *Tnf* in peritoneal leukocytes and increased the abundance of IL-1B, TNF-α, IL-6, and CXCL1 in the peritoneal lavage fluid (Figure 3f&g). Taken together, the results demonstrate that CoA, in addition to enhancing the IL-4 response, elicits a pro-inflammatory response *in vitro* and *in vivo*.

### CoA is a weak TLR4 agonist

We then hypothesized that CoA could act as an agonist for a specific TLR. Since CoA stimulated the expression of multiple pro-inflammatory genes linked to MyD88 – a signaling adaptor protein necessary for full activation of several murine toll-like receptors^37^ (Fig. 3c) – we examined whether the pro-inflammatory response from CoA could persist in BMDMs lacking MyD88. BMDMs harvested from *Myd88*^-/-^ mice significantly reduced the expression of pro-inflammatory genes induced by CoA, but marginal expression of *Il1b*, *Tnfa*, and *Irg1* persisted (Fig. 4a). The result suggested both MyD88-dependent and -independent signaling cascades underlie the pro-inflammatory response.

**Figure 4.**
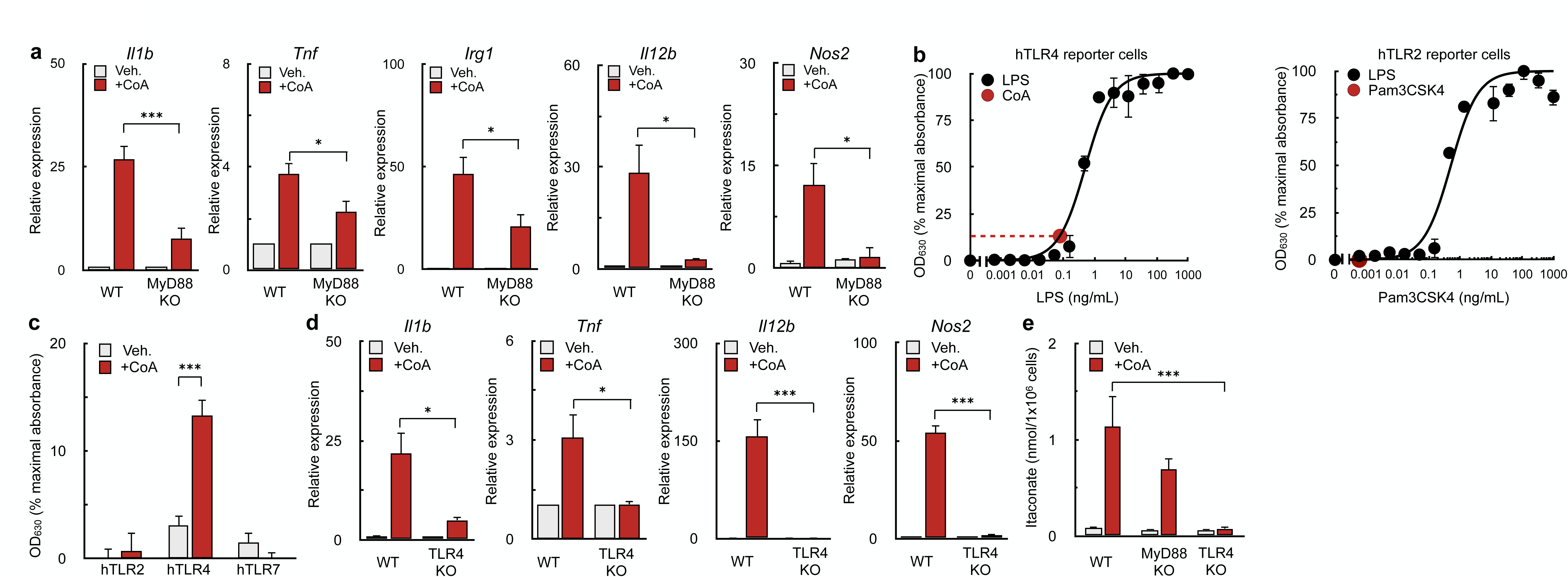
CoA is a TLR4 agonist. **(a)** qPCR analysis of *Il1b*, *Tnf*, *Irg1*, *l12b,* and *Nos2* in WT and *Myd88*^-/-^ BMDMs stimulated with 1 mM CoA or vehicle control for 4 hr. (n≥3 independent biological replicates). **(b)** Concentration-response curve of linked alkaline phosphatase activity in hTLR4 reporter cells with varying concentrations of LPS (black dots). The red dot and dashed lines represent the OD_630_ observed in response to 1 mM CoA treatment (n=4 independent biological replicates). **(c)** Aggregated response of 1 mM CoA compared to vehicle control in hTLR2, hTLR4, and hTLR7 relative to maximum TLR activation. (n≥3 independent biological replicates). **(d)** qPCR analysis of *Il1b*, *Tnf*, *l12b,* and *Nos2* in WT and *Tlr4*^-/-^ BMDMs following treatment with 1 mM CoA for 4 hr. (n=3 independent biological replicates). **(e)** Abundance of itaconate in WT, *Myd88*^-/-^, and *Tlr4*^-/-^ BMDMs in response to 1 mM CoA treatment for 48 hr. or vehicle control (n≥5 independent biological replicates). All data are presented as mean ± SEM. *p < 0.05; ***p < 0.001.

We therefore hypothesized that CoA was a TLR4 agonist. Toll-like receptor 4 (TLR4) elicits its inflammatory response by activating both MyD88-dependent and -independent signaling arms^38^. The MyD88-dependent pathway of TLR4 causes increased expression of cytokines such as *Il1b*^39^, while the TIR-domain-containing adapter-inducing interferon-β (TRIF) pathway is independent of MyD88 and increases expression of ISGs and production of type 1 interferons^40,41^. Indeed, previous results showed CoA supplementation stimulated expression of the ISG *Mx1* (Fig. 3d).

To determine if CoA is a TLR4 agonist, we utilized a reporter cell line which secretes alkaline phosphatase in response to TLR agonism^42^. Addition of CoA activated a cell line expressing human TLR4 (hTLR4) (Fig. 4b, c) but no effect was observed in cells expressing hTLR2 or hTLR7, other Myd88-linked TLRs (Fig. 4c, S3a). Interpolation of an LPS standard curve showed that 1mM CoA had a comparable effect to 0.1ng/mL LPS (Fig. 4b), indicating CoA is a relatively weak TLR4 agonist. BMDMs harvested from *Tlr4^-/-^* mice further confirmed that CoA acts via TLR4, as CoA did not increase expression of pro-inflammatory genes (Fig. 4d) or production or itaconate (Fig. 4e) in TLR4-deficient macrophages.

It was next essential to confirm that the pro-inflammatory response was due to CoA itself rather than an impurity from the >85% pure, yeast-derived CoA used in this study. In support of a direct effect of CoA, cells treated with 99% pure, synthetically-derived CoA elicited a more potent pro-inflammatory response relative to biologically-derived CoA (Fig. S3b,c). Furthermore, both the yeast-derived and synthetically-derived CoA were free of endotoxin as determined by a Limulus test (Fig. S3d). In total, the data demonstrate that CoA directly induces an inflammatory response by acting as a weak TLR4 agonist.

### Myd88-linked TLR agonists enhance the IL-4 response

Next, we asked which signaling cascade downstream of TLR4 was mediating the enhanced AAM differentiation. Although previous studies have shown that exposure to LPS and interferon gamma (IFN-γ) inhibits the acquisition of m(IL-4)^26,43^, much of this work has used high concentrations of LPS that correspond to effects that are orders of magnitude greater than that of CoA (calibrated to 0.1 ng/mL; Fig. 4b) and activate both MyD88 and TRIF.

We therefore activated macrophages with IL-4 and other TLR ligands that activate either MyD88- or TRIF-dependent signaling. Co-treatment with the MyD88-linked TLR2 agonist Pam3CSK4 (Pam3) increased the population of CD206^+^/CD301^+^ BMDMs, whereas this population was decreased upon co-treatment with the TRIF-linked TLR3 agonist Poly (I:C) (Figs. 5a&b). We also stimulated cells with IL-4 in combination with 0.1ng/mL LPS, and indeed observed increased expression of IL-4 dependent cell-surface markers (Figs. 5a&b). The TLR5 agonist flagellin and the TLR7 agonist imiquimod, both of which are upstream of MyD88, also increased expression of these IL-4-linked cell surface markers.

**Figure 5.**
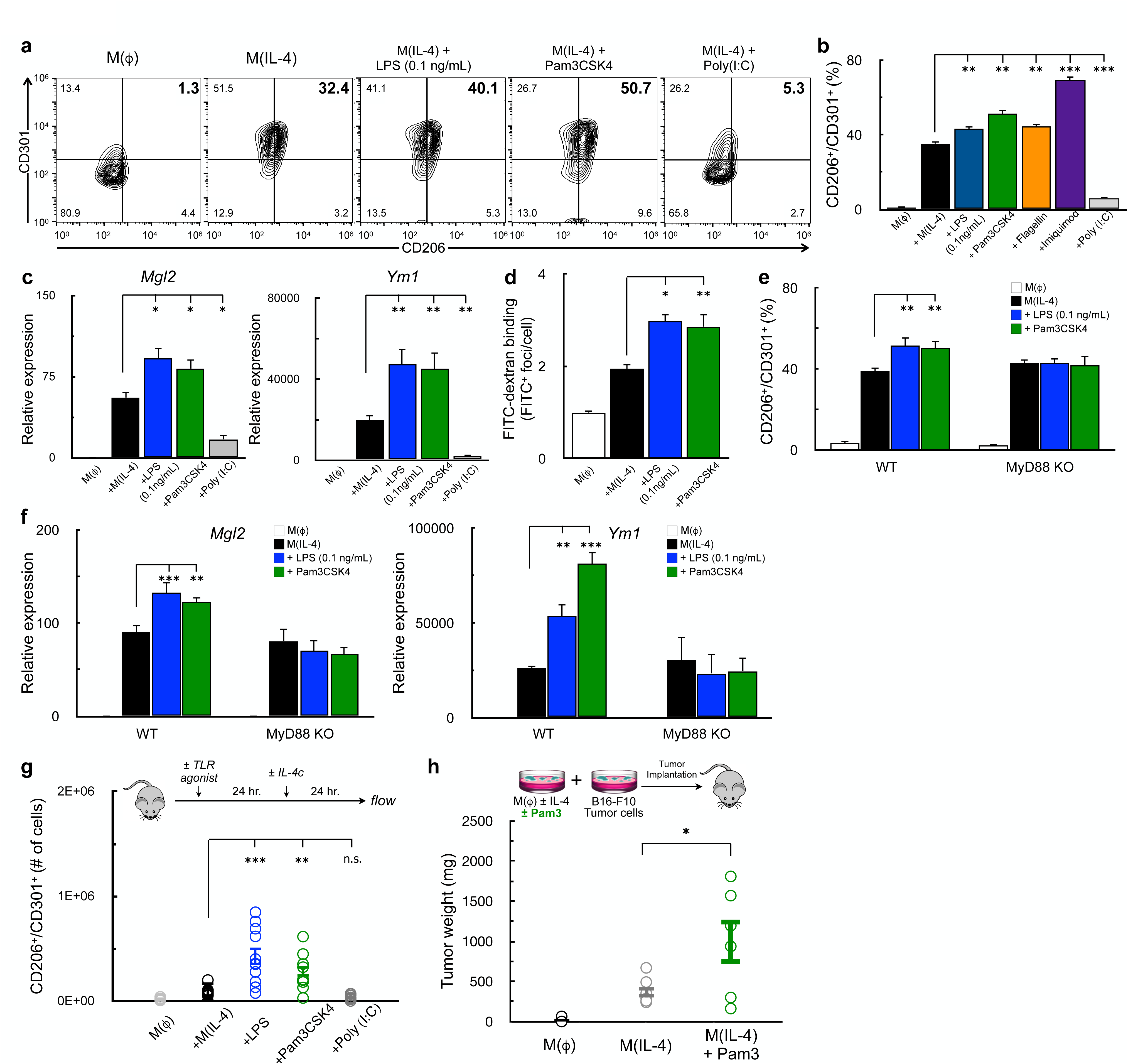
Myd88-linked TLR-ligands enhance IL-4 the response. **(a-b)** Flow cytometric analysis of CD206 and CD301 in BMDMs stimulated for 48 hr. with vehicle control, IL-4 (20 ng/mL), or IL-4 in combination with one of the following: LPS (0.1 ng/mL), Pam3CSK4 (5ng/mL), Imiquimod (10 µM), Flagellin (100 ng/mL), or Poly (I:C) (1 µg/mL) (a) Contour plots showing the percentage of cells expressing both CD206 and CD301 in the upper right quadrant. Data are from a single representative experiment. (b) Percentage of CD206^+^/CD301^+^ populations for treatments as in (a) (n=8 independent biological replicates). **(c)** qPCR analysis of *Mgl2* and *Ym1* in BMDMs stimulated with vehicle control, IL-4, or IL-4 in combination with LPS, Pam3CSK4, or Poly (I:C) for 48h. Concentrations as in (a) (n≥3 independent biological replicates). **(d)** Aggregate FITC^+^ foci per cell for BMDMs stained with FITC-Dextran and treated as with vehicle control, IL-4, or IL-4 with either LPS or Pam3CSK4 as in (a). (n=4 independent biological replicates). **(e)** Percentage of CD206^+^/CD301^+^ populations in WT and *Myd88*^-/-^BMDMs stimulated with vehicle control, IL-4, or IL-4 with either LPS or Pam3CSK4 as in (a) (n=8 independent biological replicates. **(f)** qPCR analysis of *Mgl2* and *Ym1* for BMDMs as in (e) (n=4 independent biological replicates). **(g)** Quantification of CD206^+^/CD71^+^ peritoneal macrophages from mice that were injected with vehicle control, IL-4c (5 µg IL-4 and 25 µg anti-IL-4 monoclonal antibody), IL-4c + LPS (125 µg), IL-4c +Pam3CSK4 (50 µg), or IL-4c + Poly (I:C) (200 µg). (n≥9 mice were used for each group). **(h)** Weights of subcutaneous B16-F10 melanoma tumors that were derived from the co-implantation of B16-F10 tumor cells and BMDMs that were stimulated with either vehicle control, IL-4, or IL-4 in combination with Pam3CSK4. All data are presented as mean ± SEM. *p < 0.05; **p < 0.01; ***p < 0.001; ns, not significant.

Indeed, LPS (0.1 ng/mL) and Pam3 also increased expression of IL-4-associated genes (Fig. 5c) and mannose receptor activity (Fig. 5d). As with cell surface marker expression, Poly (I:C) co-treatment lowered the expression of IL-4-stimulated genes. To definitively show that MyD88-linked signaling could affect AAM differentiation, we examined whether a low concentration of LPS or Pam3 could enhance the IL-4 response in the absence of MyD88. As expected, the effect of LPS and Pam3 on IL-4-associated cell-surface markers (Fig. 5e) and genes (Fig. 5f) was lost in BMDMs isolated from *Myd88*^-/-^ mice. Unexpectedly, both CoA and imiquimod still enhanced the IL-4 response even in the absence of MyD88 (Fig. S4). However, both compounds contain adenine-like moieties, and may interact with additional plasma membrane receptors that affect the IL-4 response independently of MyD88-linked signaling.

Lastly, we determined whether MyD88 agonists could improve M(IL-4) *in vivo* using two independent approaches. First, intraperitoneal injections of either 125µg of LPS or 25µg Pam3 prior to IL-4 complex increased the number of CD206^+^/CD301^+^ cells harvested from the peritoneal cavity, whereas no difference was observed with Poly I:C. (Fig. 5g). As further proof-of-concept, we leveraged a tumor model where alternative macrophage activation supports the growth of implanted B16 melanoma tumors^15,44^. In line with our previous results, co-treatment of IL-4-stimulated BMDMs with Pam3 resulted in significantly larger tumors when mixed with B16 melanoma cells relative to IL-4 alone (Fig. 5h). In total, these results indicate that activation of the MyD88 pathway enhances alternative macrophage activation *in vitro* and *in vivo*.

### MyD88 alters the chromatin accessibility of alternatively activated macrophages

Recent studies have highlighted the critical role of epigenetic remodeling when macrophages are exposed to a mix of pro- and anti-inflammatory ligands^45,46^. We therefore hypothesized that one mechanism by which MyD88 signaling could augment m(IL-4) was by increasing chromatin accessibility in the promoter regions of IL-4 target genes. To test this, we conducted an Assay for Transposase-Accessible Chromatin with high-throughput sequencing (ATAC-seq) analysis on IL-4-stimulated BMDMs with or without co-stimulation with Pam3. We first generated a list of the 10,878 genomic regions which had increased accessibility following IL-4 stimulation [log_2_ fold change (LFC) >1, false discovery rate (FDR) <0.01] relative to vehicle controls. These regions were localized mainly in intergenic and intronic regions (Fig. 6a, S5a).

**Figure 6.**
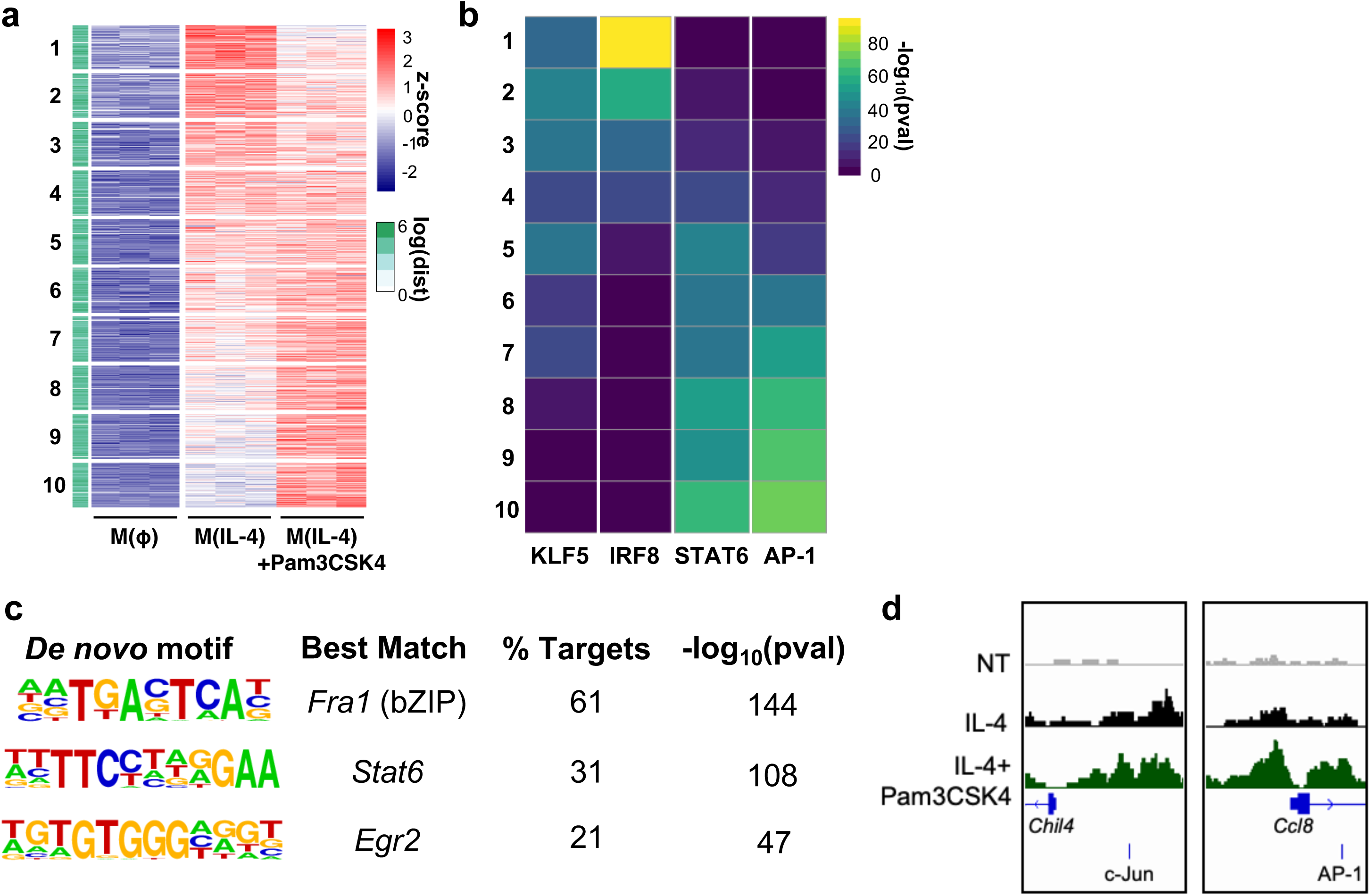
MyD88-linked signaling increases chromatin accessibility in IL-4-stumulated macrophages. **(a)** Heatmap of z-scored ATAC-seq signal for the 10,878 IL-4-inducible regions. The heatmap is arranged by increasing values for IL-4 with Pam3CSK4 co-treatment group and is divided into 10 equal bins. Side bar indicates distance to closest transcription start site (TSS). **(b)** Heatmap showing p-values of the most highly enriched motifs for each of the 10 bins that were generated in (a). **(c)** Top 3 hits from de novo transcription factor motif analysis on the significantly induced regions by Pam3CSK4 co-treatment (log_2_ fold change >0.5, false discovery rate < 0.05). **(d)** Representative tracks of Chil4 and Ccl8 promoter regions with nearby c-Jun/AP-1 motifs.

We next assessed whether these IL-4-induced regions had increased accessibility upon co-treatment with Pam3 by creating 10 equal bins in increasing order of LFC values (Fig. 6a, S5b). The analysis revealed over 30% of the IL-4-induced regions were more accessible with TLR2 co-treatment (Bins 8-10; Fig. 6b). To identify potential transcription factors that may mediate the ability of MyD88 to increase alternative activation, we then conducted HOMER transcription factor motif analysis^47^. The analysis indicated that these IL-4-induced regions with increased accessibility following Pam3 co-treatment (Bins 8-10) were enriched for STAT6 and Jun/AP-1 binding motifs (Fig. 6b). Moreover, when we expanded our analysis to consider all regions significantly increased by Pam3 (LFC >0.5 and FDR <.05, n = 1766), we noted that these regions were enriched with motifs for the AP-1 subunit *Fra1*, *Stat6* and *Egr2*. (Fig. 6c). Consistent with our hypothesis, the promoter regions of *Chil4* and *Ccl8,* genes we previously associated with the IL-4 response (Fig. 1a), were significantly more accessible upon exposure to Pam3 (Fig. 6d). Additionally, both had significantly more accessible Jun/AP-1 binding motifs following Pam3 co-treatment. Other genes associated with alternative activation such as *Pdcd1gl2* and *Arg1* also had consistent, though not statistically significant, increases in accessibility of their promoter regions following Pam3 co-treatment (Fig. S5c). In total, ATAC-Seq analysis shows that MyD88 activation regulates chromatin accessibility in AAMs. Further, the data identify the Jun/AP-1 family of transcription factors as candidates that may mediate the synergy between the MyD88 pathway and the IL-4 response.

## DISCUSSION

Our results demonstrate that the ubiquitous metabolic cofactor CoA enhances m(IL-4) in *in vitro* and *in vivo*. Genetic and pharmacologic proof-of-concept studies show, surprisingly, that CoA is a TLR4 agonist and augments alternative activation via MyD88-linked signaling. This discovery and associated data have implications for the metabolic instruction of alternative macrophage activation, the plasticity of macrophage polarization, and the breadth of intracellular metabolites and co-factors that can act as DAMPs.

Unexpectedly, addition of exogenous CoA did not augment alternative activation by increasing flux through metabolic pathways linked with m(IL-4). In fact, hallmarks of the IL-4 response such as increased respiratory capacity^17,26,27^ and *de novo* lipid synthesis^31^ were significantly reduced by CoA provision. In fact, other work using exogenously added prostaglandins shows AAM markers can further increase beyond what is induced by IL-4 while simultaneously decreasing mitochondrial oxidative metabolism^17^. The result suggests there is flexibility in the metabolic phenotypes that can support alternative macrophage activation, and provides an additional data point to the mixed results regarding whether healthy oxidative phosphorylation is obligatory for m(IL-4)^13,27^.

Additionally, gain- and loss-of-function experiments with chemical modulators of intracellular CoA levels revealed that altering CoA levels do not adjust alternative macrophage activation. The rationale for examining this hypothesis arose from studying the effects of the CPT-1 inhibitor etomoxir. High, off-target concentrations of the drug block the macrophage IL-4 response and deplete intracellular CoA, and both phenotypes were rescued upon addition of exogenous CoA^13^. The findings presented here, however, show the link between intracellular CoA levels and alternative macrophage activation is likely associative, and suggest the effects of high concentrations of etomoxir are rescued by MyD88-linked signaling rather than restoration of steady-state CoA levels. Although several independent lines of evidence show that etomoxir blocks the IL-4 response via a mechanism independent of fatty acid oxidation^13–15,48,49^ it remains unclear why high concentrations of this lipophilic, reactive epoxide block alternative activation.

Various cellular metabolites, including lipids, ATP, and uric acid can function as DAMPs to elicit an *in vitro* pro-inflammatory response^50–52^. Others, such as the complex lipid prostaglandin E2 and adenosine, can enhance alternative activation^17,53^. Here we show that CoA is a putative DAMP that primes macrophages for alternative activation and supports resolution via extrinsic, weak agonism of pro-inflammatory TLR4 and MyD88 signaling.

This aligns with recent reports that show CoA increases the expression of pro-inflammatory genes such as *Il1b*, *Tnf* and *Nos2* in mouse and human macrophages, an effect lost in mice with simultaneous genetic ablation of TLR2, TLR4, and the TLR chaperone protein Uncb93b1^54^. The data presented here that CoA itself is a TLR4 agonist likely explains the effect, rather than CoA having an indirect effect on the pro-inflammatory response via altered mitochondrial metabolism. As CoA consists of an adenosine diphosphate group linked to a phosphopantetheine moiety^32^, CoA is chemically distinct from many well characterized TLR4 agonists^55^. Interestingly, nucleoside analogues such as imiquimod (a guanosine analogue) and CL264 (an adenine analogue) are potent TLR7/8 agonists. However, CoA did not activate hTLR7 reporter cells, and additional work is required to understand the structural specificity that enables CoA to specifically activate TLR4.

TLR4 is well characterized for its ability to respond to ligands that are derived from microbes, and its role in sensing endogenous ligands is increasingly appreciated^56^. Release of intracellular proteins including tenascin 1 and high-mobility group box 1 (HMGB1) can induce a TLR4-dependent inflammatory response^42,57,58^, but the capacity for intracellular metabolites and metabolic co-factors to activate TLR4 is less established^59^. Here we show with genetic and pharmacologic proof-of-concept studies that CoA is an endogenous metabolic co-factor that can extrinsically activate TLR4 at physiologically relevant concentrations. Intracellular CoA concentrations can reach over 100 µM in the cytosol and up to 5 mM in the mitochondrial matrix^32,34,60^. As such, cell injury or death could release sufficient CoA to trigger a TLR4-mediated inflammatory response in nearby innate immune cells, particularly in regions where DAMPs can accumulate such as poorly vascularized areas. Other intracellular metabolites function as DAMPs following their cell death-induced release^61,62^, and given that CoA contains an ADP moiety, the findings are broadly consistent with the finding that adenosine can enhance the macrophage IL-4 response via the A2 adenosine receptor^53^.

Although macrophage polarization is often bifurcated into classical or alternative activation, the physiological and pathological induction of an innate immune response involves heterogeneity in activation signals and functional state^3,63,64^. For example, healing processes such as muscle and skin repair are characterized by both an initial influx of pro-inflammatory macrophages to stem infection, as well as a subsequent increase in the presence of alternatively activated macrophages to promote resolution and tissue repair^65,66^. This plasticity can be important for proper function. For example, inhibition of the initial pro-inflammatory response dampens the future expression of alternative activation markers and decreases wound healing^67^.

Cooperativity between classical and alternative macrophage activation has been established for more than 20 years, with studies demonstrating that IL-4 can enhance the expression of LPS-induced pro-inflammatory genes ^68^. Recent studies have corroborated these findings by demonstrating that IL-4 epigenetically primes macrophages to enhance the transcriptional response to pro-inflammatory stimuli^46^. Indeed, our work suggests that pro-inflammatory MyD88-linked signaling enhances the IL-4 response, at least in part, by altering chromatin accessibility.

Recent work provides strong support that MyD88-linked signaling via TLR4 agonism is a plausible mechanism by which CoA can augment m(IL-4)^69–72^. For example, the fungal effector protein CLP-1, a TLR4 agonist, fails to enhance alternative activation in MyD88^-/-^ BMDMs. Moreover, other work has associated AAMs with IL-33^73^, which is upstream of MyD88. Others have also shown that interferon-β decreases the expression of cell surface markers and genes associated with alternative activation^45,74,75^, further supporting our findings that TLR4 agonism by CoA enhances m(IL-4) via MyD88 rather than TRIF. The ATAC-Seq analysis presented here furthers this work by identifying Jun/AP-1 signaling as a candidate pathway downstream of MyD88 that may mediate the enhanced IL-4 response.

## MATERIALS AND METHODS

### Animals

All animal protocols and procedures were approved and performed in accordance with the NIH Guide for the Care and Use of Laboratory Animals and the UCLA Animal Research Committee (ARC).

### Reagents

Unless otherwise specified, yeast-derived CoA that was purchased from Sigma-Aldrich (C4780; ≥85% purity) was used. Synthetic CoA was purchased from Avanti Polar Lipids (870700P; >99% purity). IL-4 was purchased from PeproTech (214-14), and IL-4-reactive monoclonal antibodies for *in vivo* studies (IL-4 MAb) were purchased from BioXCell (BE0045). All TLR agonists were purchased from Invivogen: Pam3CSK4 (TLR2 ligand; tlrl-pms), Poly(I:C) (TLR3 ligand; tlrl-pic), LPS (TLR4 ligand; tlrl-smlps), Flagellin (TLR5 ligand; tlrl-stfla), and imiquimod (TLR7 agonist; tlrl-imqs-1).

### Isolation and differentiation of mouse BMDMs

BMDMs from wild-type (000664), *Myd88*^-/-^ (009088), and *Tlr4*^-/-^ (029015) mice were isolated as previously described from male mice aged between 6-12 weeks ^76^. Briefly, bone marrow cells were first isolated by flushing the femurs and tibiae with phosphate buffered saline (PBS). Cells were then pelleted by centrifugation at 365g for 7 mins at room temperature. Following the removal of red blood cells with RBC Lysis Buffer (Sigma, R7757) and centrifugation, bone marrow cells were differentiated for 6 days at 37°C in a humidified 5% CO_2_ incubator in ‘BMDM differentiation medium’. BMDM differentiation medium consisted of DMEM (Gibco #11965) supplemented with 10% (v/v) fetal bovine serum (FBS; Hyclone), 2mM L-glutamine, 500µM sodium pyruvate, 100 units/mL, 100 mg/mL penicillin/streptomycin,. Additionally, the medium was further supplemented 10% (v/v) with the conditioned medium from CMG-14-12 cells as a source of macrophage colony stimulate factor (M-CSF).

### In vitro BMDM activation and stimulation of Toll-like receptors

After six days of differentiation, BMDMs were scraped with a cell lifter, counted, and replated at the listed cell densities into the relevant assay format (e.g., six-well tissue culture dish, Seahorse XF96-well plate, etc.) in differentiation medium. After two days, cells were stimulated with compounds as indicated below and in the figure legends. Unless otherwise specified, all measurements of the macrophage IL-4 response were conducted 48 hr. after treatment. When assessing the effect of CoA on the pro-inflammatory response, BMDMs were treated for 4 hr. (gene expression) or 24 hr. (itaconate abundance). Unless otherwise indicated in the figures and legends, effector compounds were used at the following concentrations alongside matched vehicle controls: IL-4 (20 ng/mL), CoA (1 mM), Pam3CSK4 (5 ng/mL), Poly(I:C) (1µg/mL), LPS (0.1 ng/mL), flagellin (100ng/mL), and imiquimod (10µM).

### In vivo activation of peritoneal macrophages

To induce alternative activation *in vivo*, mice were treated with an IL-4 complex (IL-4c) consisting of a 2:1 molar ratio of IL-4 and anti-IL-4 mAb or PBS as a control. Each IL-4c-treated mouse received 5μg IL-4 and 25μg of anti-IL-4 mAb. To test the effect of CoA on alternative activation *in vivo* (Fig. 1g), mice were pretreated with an injection of either PBS or 40 mg/kg CoA for 6 hr. prior to IL-4c administration according to the scheme in the figure. After 24 hr. peritoneal macrophages were collected by rinsing the peritoneal cavity with 5mL PBS, and alternative activation markers were assessed using flow cytometry.

To determine if CoA could induce a pro-inflammatory response *in vivo* (Figs. 3f&g), mice were treated with either PBS or 40 mg/kg CoA for 6 hr. After treatment, peritoneal macrophages were collected by rinsing the peritoneal cavity with 5mL of PBS. Cells were then pelleted, with supernatant used to measure cytokine levels with the LEGENDplex multiplex ELISA kit (Biolegend, 740848), while gene expression was measured in the peritoneal exudate cells.

To determine if TLR agonists could enhance the IL-4 response of peritoneal macrophages *in vivo* (Fig. 5g), Mice were treated with either PBS or the indicated TLR agonist at the following doses: Pam3CSK4 (50μg), Poly:IC (200μg), or LPS (125μg). After 24 hr. IL-4c was administered as before, and the number of cells double-positive for CD206 and CD301 was assessed by flow cytometry.

### B16 melanoma growth

Prior to *in vivo* co-injection, *in vitro* BMDMs were either stimulated with vehicle control, IL-4, or IL-4 in combination with Pam3CSK4 for 48 hrs. On the day of implantation, a 1:1 mixture of 1×10^5^ B16-F10 cells and 1×10^5^ BMDMs were suspended in PBS and injected into the rear right flanks of 12 week old Male C57BL/6 mice^15^. Mice were sacrificed 20 days post injection and subcutaneous tumors were excised, blotted dry, and weighed.

### Gene expression analysis

Gene transcript levels were measured using qPCR. On Day 6 of differentiation, BMDMs were seeded at 3.0 ×10^5^ cells/well in 12-well plates. After activating cells with the concentrations and durations as described earlier, RNA was extracted using the RNeasy Mini Kit (Qiagen, 74106) and cDNA was synthesized using high-capacity cDNA reverse transcription kit (Applied Biosystems, 4368814) according to the manufacturers’ protocol. qPCR was performed with PowerUp SYBR green master mix (Applied Biosystems, A25743) on a QuantStudio 5 RT-PCR (Applied Biosystems). Relative gene expression values were calculated using the delta-delta Ct method, with the ribosomal protein *Rplp0* used as a control for normalization.

### Flow cytometric analysis

Cell surface marker expression was measured using flow cytometry. On Day 6 of differentiation, BMDMs were seeded at 3.0 ×10^5^ cells/well in 12-well plates. After activating cells with the concentrations and durations as described earlier, BMDMs were detached by scraping in 450µL of Accutase. Cells were then washed with FACS buffer (PBS+ 2% (v/v) FBS with 1mM EDTA) and incubated with a 1:500 dilution of TruStain FCX (Biolegend, 101320) for 5 minutes. Cells were next stained for 30 mins on ice with a 1:300 dilution of antibodies raised against mouse CD206 (Biolegend,141710), CD301(Biorad, MCA2392A647T), or CD71(Biolegend, 113812). Cells were then washed and resuspended in FACS buffer containing 1 μg/mL DAPI (Invitrogen, D1306) for viability analysis, and data was captured on an Attune NXT flow cytometer. Data were analyzed using FlowJo X software.

### Endocytosis assay

Following differentiation, BMDMs were seeded in black-walled 96-well plates at 3 ×10^4^ cells/well for high-content imaging to quantify uptake of FITC-dextran. 48 hr. after compound treatment, medium was replaced with high-glucose DMEM containing 1mg/mL FITC-dextran (Sigma, FD40) and 10ng/mL Hoechst 33342. Following a 1hr. incubation at 37°C, cells were washed twice with PBS and fixed with 4% (v/v) paraformaldehyde (PFA) in PBS. Images were captured with a PerkinElmer Operetta, and FITC-positive foci per cell was calculated using Harmony software.

### Quantification of short-chain acyl CoAs

Quantification of acyl CoAs was conducted according to previously established methods^18^. Following differentiation, BMDMs were seeded in 10cm^2^ dishes at 5 ×10^6^ cells/dish. Following 48hr. stimulation, cells were rinsed twice with ice-cold PBS, scraped into 1.5mL microfuge tubes and pelleted via centrifugation at 4°C. 200μL of an ice-cold extraction solution [2.5% (w/v) 5-sulfosalicylic acid (SSA) along with 1μM Crotonoyl CoA as an internal standard] was added to each cell pellet and subsequently vortexed. Samples were centrifuged at 18,000*g* for 15 min at 4°C. Supernatants containing short-chain acyl CoAs were then removed and transferred to glass LC-MS vials for analysis as is thoroughly described elsewhere^18^.

### Seahorse XF Analysis

After 6 days of differentiation, BMDMs were plated at 3.0 x 10^4^ cells/well in XF96 plates. Following 48 hr. of treatment with compounds under investigation, respirometry assays were conducted with an Agilent Seahorse XFe96 Analyzer. Oligomycin (2 μM), two injections of FCCP (750 nM each), and rotenone (200 nM) with antimycin A (1 μM) were added acutely to the wells, and respiratory parameters calculated according to best practices^77,78^. Measurements were conducted in unbuffered DMEM (Sigma #5030) supplemented with 5 mM HEPES, 8 mM glucose, 2 mM glutamine, and 2 mM pyruvate. Lactate efflux was measured by correcting rates of extracellular acidification for microplate sensor coverage and confounding respiratory acidification^79^.

#### Metabolomics and stable isotope tracing of polar metabolites

After 6 days of differentiation, BMDMs were plated at 1 ×10^6^ cells/well in 6-well dishes. Cells were then stimulated in medium where either glucose or glutamine was replaced with uniformly labeled ^13^C_6_-glucose (CLM-1396) or uniformly labeled ^13^C_5_-glutamine (CLM-1822). After 48 hr., cells were harvested and extracted for GC/MS using established methods, with all steps conducted on ice^80^. Briefly, cell plates were washed twice with 0.9% (w/v) NaCl and samples were extracted with 500 μL methanol, 200 μL water containing 1 μg of norvaline (internal standard), and 500 μL chloroform. Samples were vortexed for 1 min and spun at 10,000 *g* for 5 min at 4°C, and the aqueous layers containing the polar metabolites were transferred to GC/MS sample vials and dried overnight using a refrigerated CentriVap. Once dry, the samples were resuspended in 20 μL of 2% (w/v) methoxyamine in pyridine and incubated at 37°C for 45 minutes. This was followed by addition of 20 μL of MTBSTFA + 1% TBDMS (N-tert-Butyldimethylsilyl-N-methyltrifluoroacetamide with 1% tertButyldimethylchlorosilane). Following a second 45-minute incubation at 37°C, samples were run as previously described ^80^. Analysis was conducted using Agilent MassHunter software, and stable isotope tracing data was corrected for natural abundance of heavy isotopes with FluxFix software using a reference set of unlabeled metabolite standards ^81^.

### De novo lipogenesis

Briefly, after 6 days of differentiation, BMDMs were plated at 1 ×10^5^ cells/ well in 24-well dishes Cells were stimulated in medium in which unlabeled glucose was replaced with 10mM uniformly labeled ^13^C_6_-glucose. Extraction of fatty acids, quantification of *de novo* synthesis, and normalization to cell number was conducted using an Agilent 5975C mass spectrometer coupled to a 7890 gas chromatograph as previously described^76,82^.

### HEK-Blue hTLR reporter assays

HEK-Blue reporter cells expressing either human TLR2 (hkb-htlr2), hTLR4 (hkb-htlr4), or hTLR7 (hkb-htlr7v2) were purchased from InvivoGen and maintained according to the manufacturer’s instructions. To establish concentration-response curves, 2.5 x 10^3^ reporter cells were resuspended in HEK-Blue detection medium (Invivogen, hb-det2) in 96-well plates and stimulated with the appropriate agonist (Pam3CSK4: 0.6pg/mL to 1 µg/mL; LPS: 0.6pg/mL to 1 µg/mL; CL307: 0.6pg/mL to 1 µg/mL). Following a 24hr. incubation, secreted alkaline phosphatase reporter activity was determined by assessing OD_630_ with a plate reader. The OD_630_ of cells treated with 1 mM CoA was compared relative to positive controls. Calibration cuves to fit empirical data were generated using GraphPad Prism software.

### ATAC-Seq library prep

ATAC-seq libraries were produced by the Applied Genomics, Computation and Translational Core Facility at Cedars Sinai. Briefly 50,000 BMDMs per sample were lysed to collect nuclei and treated with Tn5 transposase (Illumina) for 30 min at 37°C with gentle agitation. The DNA was isolated with DNA Clean & Concentrator Kit (Zymo) and PCR amplified and barcoded with NEBNext High-Fidelity PCR Mix (New England Biolabs) and unique dual indexes (Illumina). The ATAC-seq library amplification was confirmed by real-time PCR, and additional barcoding PCR cycles were added as necessary while avoiding overamplification. Amplified ATAC-seq libraries were purified with DNA Clean & Concentrator Kit (Zymo). The purified libraries were quantified with Kapa Library Quant Kit (KAPA Biosystems) and quality assessed on a 4200 TapeStation System (Agilent). The libraries were pooled based on molar concentrations and sequenced on a HiSeq 4000 platform (paired end, 100 bp).

### ATAC-Seq analysis

The peaks for all the ATAC-seq samples were used to generate a single reference peak file, and the number of reads that fell into each peak was counted using deeptools multiBamSummary^83^. EdgeR^84^ was used to determine the IL-4 significantly induced regions by applying a cutoff FDR <0.01 and LFC > 1 of triplicate data upon IL-4 stimulation on WT BMDMs and to determine the Pam3CSK4 co-treatment significant regions by applying a cutoff FDR <0.05 and LFC > 0.5. Analysis of transcription factor motif enrichment was performed using findMotifsGenome function in the HOMER suite^47^, using all detected peaks as background. Reads were normalized by RPKM. Data were visualized with ggplot2 or the pheatmap packages in R.

### RNA-Seq library prep and quantification

RNA sequencing libraries were produced by the Technology Center for Genomics & Bioinformatics at UCLA. Isolation of RNA was performed using Qiagen RNeasy Mini kit and RNA libraries were prepared with KAPA stranded mRNA-Seq kit. High throughput sequencing was performed on Illumina NovaSeq 6000 (paired end, 2×150bp) targeting 100 million reads per sample. Demultiplexing was performed with Illumina Bcl2fastq v2.19.1. Gene expression quantification from the resulting fastqs was performed using Salmon v1.21.1 in mapping-based mode (Patro et al. 2017). Reads were selectively aligned to the GENCODE vM25 mouse reference transcriptome with corrections for sequence-specific and GC content biases.

### RNA-Seq analysis

Raw gene count data were analyzed using the R package DESeq2 v1.22.2^85^ for library size normalization and differential expression analysis. For differential expression results, genes with adjusted p-values below 0.01 and log2 fold changes above 1 or below −1 were deemed significant. For visualization in volcano plots, log2 fold changes above 10 or below −10 were set to 10 and - 10 respectively. Gene set enrichment analysis^86^ was performed using the R package FGSEA v1.15.0^87^ based on gene lists ranked by the Wald statistic from differential expression results. Genesets corresponding to the mouse transcriptome from KEGG, REACTOME, and BIOCARTA within the Molecular Signatures Database^88^ were accessed using the R package msigdbr v7.1.1^89^ Genesets with adjusted p-values below 0.05 were deemed significant.

### Statistical analysis

All statistical parameters, including the number of biological replicates (n), can be found in the figure legends. Statistical analyses were performed using Graph Pad Prism 5 software. Data are presented as the mean ± standard deviation unless otherwise specified. Individual pairwise comparisons were performed using two-tailed Student’s t-test. For analysis involving more than two groups, data were analyzed by repeated measures ANOVA followed by Dunnett’s post-hoc multiple comparisons tests (compared against vehicle controls unless otherwise specified). Data were assumed to follow a normal distribution (no tests were performed). Values denoted as follows were considered statistically significant: *, p < 0.05; **, p < 0.01; ***, p < 0.001.

## Supporting information

Manuscript Supplement

## AUTHOR CONTRIBUTIONS

Conceptualization: AEJ, SJB, ASD; Data curation: AEJ, AR, NI, CC, NAB, ASD; Formal analysis: AEJ, AR, NI, CC, NAB, ASD; Funding acquisition: TGG, AH, SJB, ASD; Investigation: AEJ, AR, NI, CC, NAB, ABB, WYH, AS, ARB, AAC; Methodology: AEJ, CC, NAB, AS, ASD; Project administration: SJB, ASD; Resources: TGG, AH, SJB, ASD; Supervision: AEJ, SJB, ASD; Writing - original draft: AEJ, ASD; Writing - review & editing: All authors

## ACKNOWLEDGEMENTS

ASD is supported by National Institutes of Health (NIH) Grant R35GM138003, the W.M. Keck Foundation (995337), and the Agilent Early Career Professor Award. SJB is supported by the NIH Grants P01HL146358 and R01HL157710. AEJ was supported by the UCLA Tumor Cell Biology Training Program T32 CA009056. ABB is supported by the UCLA Chemistry-Biology Interface Training Grant T32GM136614. AH is supported by NIH Grant R01AI173214.

## REFERENCES

1 Glass, C. K. & Natoli, G. Molecular control of activation and priming in macrophages. Nat Immunol 17, 26–33, doi:10.1038/ni.3306 (2016).

2 Murray, P. J. Macrophage Polarization. Annu Rev Physiol 79, 541–566, doi:10.1146/annurev-physiol-022516-034339 (2017).

3 Murray, P. J. et al. Macrophage activation and polarization: nomenclature and experimental guidelines. Immunity 41, 14–20, doi:10.1016/j.immuni.2014.06.008 (2014).

4 Van den Bossche, J., O’Neill, L. A. & Menon, D. Macrophage Immunometabolism: Where Are We (Going)? Trends Immunol 38, 395–406, doi:10.1016/j.it.2017.03.001 (2017).

5 Jones, A. E. & Divakaruni, A. S. Macrophage activation as an archetype of mitochondrial repurposing. Mol Aspects Med 71, 100838, doi:10.1016/j.mam.2019.100838 (2020).

6 Wculek, S. K., Dunphy, G., Heras-Murillo, I., Mastrangelo, A. & Sancho, D. Metabolism of tissue macrophages in homeostasis and pathology. Cell Mol Immunol 19, 384–408, doi:10.1038/s41423-021-00791-9 (2022).

7 Tannahill, G. M. et al. Succinate is an inflammatory signal that induces IL-1β through HIF-1α. Nature 496, 238–242, doi:10.1038/nature11986 (2013).

8 Haschemi, A. et al. The sedoheptulose kinase CARKL directs macrophage polarization through control of glucose metabolism. Cell Metab 15, 813–826, doi:10.1016/j.cmet.2012.04.023 (2012).

9 Mills, E. L. et al. Succinate Dehydrogenase Supports Metabolic Repurposing of Mitochondria to Drive Inflammatory Macrophages. Cell 167, 457–470.e413, doi:10.1016/j.cell.2016.08.064 (2016).

10 Britt, E. C. et al. Switching to the cyclic pentose phosphate pathway powers the oxidative burst in activated neutrophils. Nat Metab 4, 389–403, doi:10.1038/s42255-022-00550-8 (2022).

11 Vats, D. et al. Oxidative metabolism and PGC-1beta attenuate macrophage-mediated inflammation. Cell Metab 4, 13–24, doi:10.1016/j.cmet.2006.05.011 (2006).

12 Huang, S. C. et al. Metabolic Reprogramming Mediated by the mTORC2-IRF4 Signaling Axis Is Essential for Macrophage Alternative Activation. Immunity 45, 817–830, doi:10.1016/j.immuni.2016.09.016 (2016).

13 Divakaruni, A. S. et al. Etomoxir Inhibits Macrophage Polarization by Disrupting CoA Homeostasis. Cell Metab 28, 490–503.e497, doi:10.1016/j.cmet.2018.06.001 (2018).

14 Van den Bossche, J. & van der Windt, G. J. W. Fatty Acid Oxidation in Macrophages and T Cells: Time for Reassessment? Cell Metab 28, 538–540, doi:10.1016/j.cmet.2018.09.018 (2018).

15 Bakker, N. v. T. et al. In macrophages fatty acid oxidation spares glutamate for use in diverse metabolic pathways required for alternative activation. bioRxiv, 2022.2004.2013.487890, doi:10.1101/2022.04.13.487890 (2022).

16 Covarrubias, A. J. et al. Akt-mTORC1 signaling regulates Acly to integrate metabolic input to control of macrophage activation. Elife 5, doi:10.7554/eLife.11612 (2016).

17 Sanin, D. E. et al. Mitochondrial Membrane Potential Regulates Nuclear Gene Expression in Macrophages Exposed to Prostaglandin E2. Immunity 49, 1021–1033.e1026, doi:10.1016/j.immuni.2018.10.011 (2018).

18 Jones, A. E. et al. A Single LC-MS/MS Analysis to Quantify CoA Biosynthetic Intermediates and Short-Chain Acyl CoAs. Metabolites 11, doi:10.3390/metabo11080468 (2021).

19 Gonzalez-Hurtado, E. et al. Loss of macrophage fatty acid oxidation does not potentiate systemic metabolic dysfunction. Am J Physiol Endocrinol Metab 312, E381–e393, doi:10.1152/ajpendo.00408.2016 (2017).

20 Liu, P. S. et al. α-ketoglutarate orchestrates macrophage activation through metabolic and epigenetic reprogramming. Nat Immunol 18, 985–994, doi:10.1038/ni.3796 (2017).

21 Zhang, X. et al. CCL8 secreted by tumor-associated macrophages promotes invasion and stemness of glioblastoma cells via ERK1/2 signaling. Lab Invest 100, 619–629, doi:10.1038/s41374-019-0345-3 (2020).

22 Webb, D. C., McKenzie, A. N. & Foster, P. S. Expression of the Ym2 lectin-binding protein is dependent on interleukin (IL)-4 and IL-13 signal transduction: identification of a novel allergy-associated protein. J Biol Chem 276, 41969–41976, doi:10.1074/jbc.M106223200 (2001).

23 van Die, I. & Cummings, R. D. The Mannose Receptor in Regulation of Helminth-Mediated Host Immunity. Front Immunol 8, 1677, doi:10.3389/fimmu.2017.01677 (2017).

24 van Liempt, E. et al. Schistosoma mansoni soluble egg antigens are internalized by human dendritic cells through multiple C-type lectins and suppress TLR-induced dendritic cell activation. Mol Immunol 44, 2605–2615, doi:10.1016/j.molimm.2006.12.012 (2007).

25 Sallusto, F., Cella, M., Danieli, C. & Lanzavecchia, A. Dendritic cells use macropinocytosis and the mannose receptor to concentrate macromolecules in the major histocompatibility complex class II compartment: downregulation by cytokines and bacterial products. J Exp Med 182, 389–400, doi:10.1084/jem.182.2.389 (1995).

26 Van den Bossche, J., et al. Mitochondrial Dysfunction Prevents Repolarization of Inflammatory Macrophages. Cell Rep 17, 684–696, doi:10.1016/j.celrep.2016.09.008 (2016).

27 Puleston, D. J. et al. Polyamines and eIF5A Hypusination Modulate Mitochondrial Respiration and Macrophage Activation. Cell Metab 30, 352–363.e358, doi:10.1016/j.cmet.2019.05.003 (2019).

28 Tan, Z. et al. Pyruvate Dehydrogenase Kinase 1 Participates in Macrophage Polarization via Regulating Glucose Metabolism. The Journal of Immunology 194, 6082–6089, doi:10.4049/jimmunol.1402469 (2015).

29 Jha, A. K. et al. Network integration of parallel metabolic and transcriptional data reveals metabolic modules that regulate macrophage polarization. Immunity 42, 419–430, doi:10.1016/j.immuni.2015.02.005 (2015).

30 Huang, S. C. et al. Cell-intrinsic lysosomal lipolysis is essential for alternative activation of macrophages. Nat Immunol 15, 846–855, doi:10.1038/ni.2956 (2014).

31 Bidault, G. et al. SREBP1-induced fatty acid synthesis depletes macrophages antioxidant defences to promote their alternative activation. Nat Metab 3, 1150–1162, doi:10.1038/s42255-021-00440-5 (2021).

32 Leonardi, R., Zhang, Y. M., Rock, C. O. & Jackowski, S. Coenzyme A: back in action. Prog Lipid Res 44, 125–153, doi:10.1016/j.plipres.2005.04.001 (2005).

33 Pietrocola, F., Galluzzi, L., Bravo-San Pedro, J. M., Madeo, F. & Kroemer, G. Acetyl coenzyme A: a central metabolite and second messenger. Cell Metab 21, 805–821, doi:10.1016/j.cmet.2015.05.014 (2015).

34 Gout, I. Coenzyme A, protein CoAlation and redox regulation in mammalian cells. Biochem Soc Trans 46, 721–728, doi:10.1042/bst20170506 (2018).

35 Steinhelper, M. E. & Olson, M. S. The effects of cyclopropane carboxylate on hepatic pyruvate metabolism. Arch Biochem Biophys 243, 80–91, doi:10.1016/0003-9861(85)90775-1 (1985).

36 Sharma, L. K. et al. A therapeutic approach to pantothenate kinase associated neurodegeneration. Nature Communications 9, 4399, doi:10.1038/s41467-018-06703-2 (2018).

37 Snyder, G. A. et al. Molecular mechanisms for the subversion of MyD88 signaling by TcpC from virulent uropathogenic *Escherichia coli*. Proceedings of the National Academy of Sciences 110, 6985-6990, doi:doi:10.1073/pnas.1215770110 (2013).

38 Fitzgerald, K. A. & Kagan, J. C. Toll-like Receptors and the Control of Immunity. Cell 180, 1044–1066, doi:10.1016/j.cell.2020.02.041 (2020).

39 Kawai, T., Adachi, O., Ogawa, T., Takeda, K. & Akira, S. Unresponsiveness of MyD88-deficient mice to endotoxin. Immunity 11, 115–122, doi:10.1016/s1074-7613(00)80086-2 (1999).

40 Kawai, T. et al. Lipopolysaccharide stimulates the MyD88-independent pathway and results in activation of IFN-regulatory factor 3 and the expression of a subset of lipopolysaccharide-inducible genes. J Immunol 167, 5887–5894, doi:10.4049/jimmunol.167.10.5887 (2001).

41 Toshchakov, V. et al. TLR4, but not TLR2, mediates IFN-beta-induced STAT1alpha/beta-dependent gene expression in macrophages. Nat Immunol 3, 392–398, doi:10.1038/ni774 (2002).

42 Sosa, R. A. et al. Disulfide High-Mobility Group Box 1 Drives Ischemia-Reperfusion Injury in Human Liver Transplantation. Hepatology 73, 1158–1175, doi:10.1002/hep.31324 (2021).

43 Khallou-Laschet, J. et al. Macrophage Plasticity in Experimental Atherosclerosis. PLOS ONE 5, e8852, doi:10.1371/journal.pone.0008852 (2010).

44 Chen, X. W. et al. CYP4A in tumor-associated macrophages promotes pre-metastatic niche formation and metastasis. Oncogene 36, 5045–5057, doi:10.1038/onc.2017.118 (2017).

45 Dichtl, S. et al. Gene-selective transcription promotes the inhibition of tissue reparative macrophages by TNF. Life Sci Alliance 5, doi:10.26508/lsa.202101315 (2022).

46 Czimmerer, Z. et al. The epigenetic state of IL-4-polarized macrophages enables inflammatory cistromic expansion and extended synergistic response to TLR ligands. Immunity 55, 2006–2026.e2006, doi:10.1016/j.immuni.2022.10.004 (2022).

47 Heinz, S. et al. Simple combinations of lineage-determining transcription factors prime cis-regulatory elements required for macrophage and B cell identities. Mol Cell 38, 576–589, doi:10.1016/j.molcel.2010.05.004 (2010).

48 Ceccarelli, S. M., Chomienne, O., Gubler, M. & Arduini, A. Carnitine palmitoyltransferase (CPT) modulators: a medicinal chemistry perspective on 35 years of research. J Med Chem 54, 3109–3152, doi:10.1021/jm100809g (2011).

49 Nomura, M. et al. Fatty acid oxidation in macrophage polarization. Nat Immunol 17, 216–217, doi:10.1038/ni.3366 (2016).

50 Kim, S. M. et al. Hyperuricemia-induced NLRP3 activation of macrophages contributes to the progression of diabetic nephropathy. Am J Physiol Renal Physiol 308, F993–f1003, doi:10.1152/ajprenal.00637.2014 (2015).

51 Wang, X., Wang, Y., Antony, V., Sun, H. & Liang, G. Metabolism-Associated Molecular Patterns (MAMPs). Trends Endocrinol Metab 31, 712–724, doi:10.1016/j.tem.2020.07.001 (2020).

52 Howell, K. W. et al. Toll-like receptor 4 mediates oxidized LDL-induced macrophage differentiation to foam cells. J Surg Res 171, e27–31, doi:10.1016/j.jss.2011.06.033 (2011).

53 Csóka, B. et al. Adenosine promotes alternative macrophage activation via A2A and A2B receptors. Faseb j 26, 376–386, doi:10.1096/fj.11-190934 (2012).

54 Timblin, G. A. et al. Coenzyme A governs proinflammatory macrophage metabolism. bioRxiv, 2022.2008.2030.505732, doi:10.1101/2022.08.30.505732 (2022).

55 Romerio, A. & Peri, F. Increasing the Chemical Variety of Small-Molecule-Based TLR4 Modulators: An Overview. Front Immunol 11, 1210, doi:10.3389/fimmu.2020.01210 (2020).

56 Rifkin, I. R., Leadbetter, E. A., Busconi, L., Viglianti, G. & Marshak-Rothstein, A. Toll-like receptors, endogenous ligands, and systemic autoimmune disease. Immunol Rev 204, 27–42, doi:10.1111/j.0105-2896.2005.00239.x (2005).

57 Midwood, K. et al. Tenascin-C is an endogenous activator of Toll-like receptor 4 that is essential for maintaining inflammation in arthritic joint disease. Nat Med 15, 774–780, doi:10.1038/nm.1987 (2009).

58 Okamura, Y. et al. The extra domain A of fibronectin activates Toll-like receptor 4. J Biol Chem 276, 10229–10233, doi:10.1074/jbc.M100099200 (2001).

59 Lancaster, G. I. et al. Evidence that TLR4 Is Not a Receptor for Saturated Fatty Acids but Mediates Lipid-Induced Inflammation by Reprogramming Macrophage Metabolism. Cell Metab 27, 1096–1110.e1095, doi:10.1016/j.cmet.2018.03.014 (2018).

60 Williamson, J. R. & Corkey, B. E. Assay of citric acid cycle intermediates and related compounds--update with tissue metabolite levels and intracellular distribution. Methods Enzymol 55, 200–222, doi:10.1016/0076-6879(79)55025-3 (1979).

61 Cruz, C. M. et al. ATP activates a reactive oxygen species-dependent oxidative stress response and secretion of proinflammatory cytokines in macrophages. J Biol Chem 282, 2871–2879, doi:10.1074/jbc.M608083200 (2007).

62 Vénéreau, E., Ceriotti, C. & Bianchi, M. E. DAMPs from Cell Death to New Life. Front Immunol 6, 422, doi:10.3389/fimmu.2015.00422 (2015).

63 Gordon, S. & Martinez, F. O. Alternative Activation of Macrophages: Mechanism and Functions. Immunity 32, 593–604, 10.1016/j.immuni.2010.05.007 (2010).

64 Ruffell, B., Affara, N. I. & Coussens, L. M. Differential macrophage programming in the tumor microenvironment. Trends Immunol 33, 119–126, doi:10.1016/j.it.2011.12.001 (2012).

65 Arnold, L. et al. Inflammatory monocytes recruited after skeletal muscle injury switch into antiinflammatory macrophages to support myogenesis. J Exp Med 204, 1057–1069, doi:10.1084/jem.20070075 (2007).

66 Hesketh, M., Sahin, K. B., West, Z. E. & Murray, R. Z. Macrophage Phenotypes Regulate Scar Formation and Chronic Wound Healing. Int J Mol Sci 18, doi:10.3390/ijms18071545 (2017).

67 Lucas, T. et al. Differential Roles of Macrophages in Diverse Phases of Skin Repair. The Journal of Immunology 184, 3964–3977, doi:10.4049/jimmunol.0903356 (2010).

68 Major, J., Fletcher, J. E. & Hamilton, T. A. IL-4 pretreatment selectively enhances cytokine and chemokine production in lipopolysaccharide-stimulated mouse peritoneal macrophages. J Immunol 168, 2456–2463, doi:10.4049/jimmunol.168.5.2456 (2002).

69 O’Brien, E. M. & Spiller, K. L. Pro-inflammatory polarization primes Macrophages to transition into a distinct M2-like phenotype in response to IL-4. J Leukoc Biol 111, 989–1000, doi:10.1002/jlb.3a0520-338r (2022).

70 Liu, S. X., Gustafson, H. H., Jackson, D. L., Pun, S. H. & Trapnell, C. Trajectory analysis quantifies transcriptional plasticity during macrophage polarization. Sci Rep 10, 12273, doi:10.1038/s41598-020-68766-w (2020).

71 Rao, A. J. et al. Revision joint replacement, wear particles, and macrophage polarization. Acta Biomater 8, 2815–2823, doi:10.1016/j.actbio.2012.03.042 (2012).

72 Dang, E. V. et al. Secreted fungal virulence effector triggers allergic inflammation via TLR4. Nature 608, 161–167, doi:10.1038/s41586-022-05005-4 (2022).

73 Faas, M. et al. IL-33-induced metabolic reprogramming controls the differentiation of alternatively activated macrophages and the resolution of inflammation. Immunity 54, 2531–2546.e2535, doi:10.1016/j.immuni.2021.09.010 (2021).

74 Kusnadi, A. et al. The Cytokine TNF Promotes Transcription Factor SREBP Activity and Binding to Inflammatory Genes to Activate Macrophages and Limit Tissue Repair. Immunity 51, 241–257.e249, doi:10.1016/j.immuni.2019.06.005 (2019).

75 Ming-Chin Lee, K., et al. Type I interferon antagonism of the JMJD3-IRF4 pathway modulates macrophage activation and polarization. Cell Rep 39, 110719, doi:10.1016/j.celrep.2022.110719 (2022).

76 Hsieh, W. Y., Williams, K. J., Su, B. & Bensinger, S. J. Profiling of mouse macrophage lipidome using direct infusion shotgun mass spectrometry. STAR Protoc 2, 100235, doi:10.1016/j.xpro.2020.100235 (2021).

77 Divakaruni, A. S., Paradyse, A., Ferrick, D. A., Murphy, A. N. & Jastroch, M. Analysis and interpretation of microplate-based oxygen consumption and pH data. Methods Enzymol 547, 309–354, doi:10.1016/b978-0-12-801415-8.00016-3 (2014).

78 Divakaruni, A. S. & Jastroch, M. A practical guide for the analysis, standardization and interpretation of oxygen consumption measurements. Nat Metab 4, 978–994, doi:10.1038/s42255-022-00619-4 (2022).

79 Desousa, B. R. et al. Calculation of ATP production rates using the Seahorse XF Analyzer. EMBO Rep 24, e56380, doi:10.15252/embr.202256380 (2023).

80 Cordes, T. & Metallo, C. M. Quantifying Intermediary Metabolism and Lipogenesis in Cultured Mammalian Cells Using Stable Isotope Tracing and Mass Spectrometry. Methods Mol Biol 1978, 219–241, doi:10.1007/978-1-4939-9236-2_14 (2019).

81 Trefely, S., Ashwell, P. & Snyder, N. W. FluxFix: automatic isotopologue normalization for metabolic tracer analysis. BMC Bioinformatics 17, 485, doi:10.1186/s12859-016-1360-7 (2016).

82 Argus, J. P. et al. Development and Application of FASA, a Model for Quantifying Fatty Acid Metabolism Using Stable Isotope Labeling. Cell Rep 25, 2919–2934.e2918, doi:10.1016/j.celrep.2018.11.041 (2018).

83 Ramírez, F. et al. deepTools2: a next generation web server for deep-sequencing data analysis. Nucleic Acids Res 44, W160–165, doi:10.1093/nar/gkw257 (2016).

84 Robinson, M. D., McCarthy, D. J. & Smyth, G. K. edgeR: a Bioconductor package for differential expression analysis of digital gene expression data. Bioinformatics 26, 139–140, doi:10.1093/bioinformatics/btp616 (2010).

85 Love, M. I., Huber, W. & Anders, S. Moderated estimation of fold change and dispersion for RNA-seq data with DESeq2. Genome Biol 15, 550, doi:10.1186/s13059-014-0550-8 (2014).

86 Subramanian, A. et al. Gene set enrichment analysis: a knowledge-based approach for interpreting genome-wide expression profiles. Proc Natl Acad Sci U S A 102, 15545–15550, doi:10.1073/pnas.0506580102 (2005).

87. Korotkevich, G., et al. Fast gene set enrichment analysis. bioRxiv, 060012, doi:10.1101/060012 (2021).

88 Liberzon, A. et al. The Molecular Signatures Database (MSigDB) hallmark gene set collection. Cell Syst 1, 417–425, doi:10.1016/j.cels.2015.12.004 (2015).

89 Dolgalev, I. Vol. R package version 7.5.1.9001 (2022).

