## Supplementary material for "The metabolic cofactor Coenzyme A enhances alternative macrophage activation via MyD88-linked signaling": Manuscript Supplement

Includes Supplemental Information Figures S1–5

**Supplemental Information Figure S1**

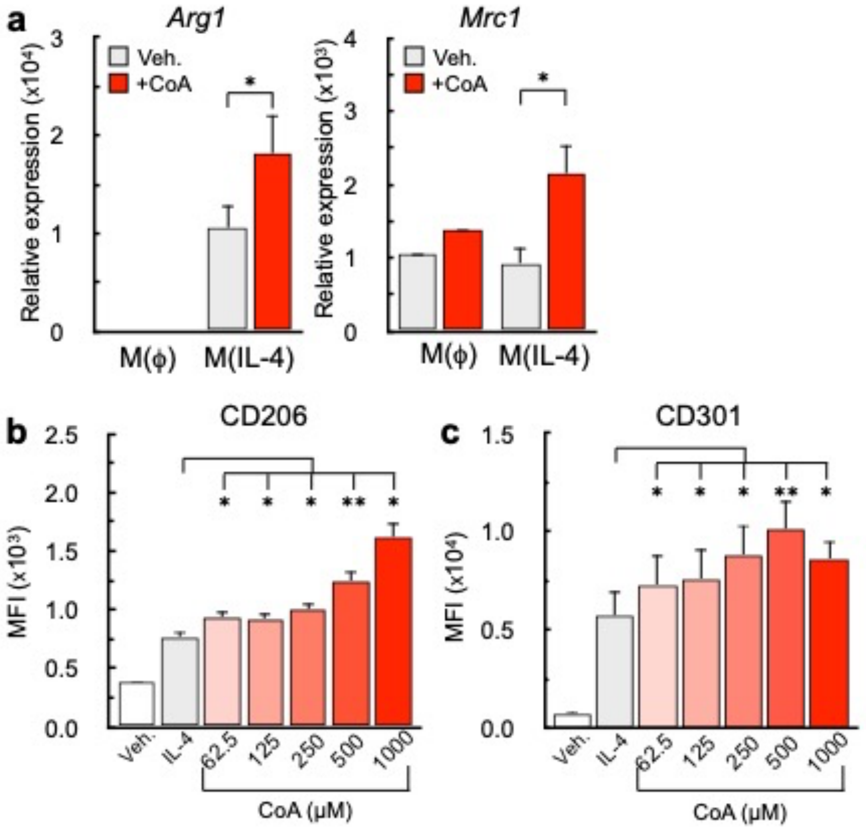


**Supplemental Figure S1. Exogenous CoA provision enhances alternative macrophage activation (a)** qPCR analysis of the IL-4-associated genes *Arg1 and Mrc1* in BMDMs treated with vehicle, CoA (1 mM), IL-4 (20 ng/mL), or CoA + IL-4 for 48 hr. (n≥9 independent biological replicates) **(b)** Mean fluorescence intensity of CD206 and CD301 in BMDMs following treatment with vehicle control, IL-4 (20 ng/mL), or IL-4 + CoA with concentrations as indicated in the figure for 48 hr. (n=4 independent biological replicates). All data are presented as mean ± SEM. $^*p < 0.05$; $^{**}p < 0.01$; $^{***}p < 0.001$.

**Supplemental Information Figure S2**

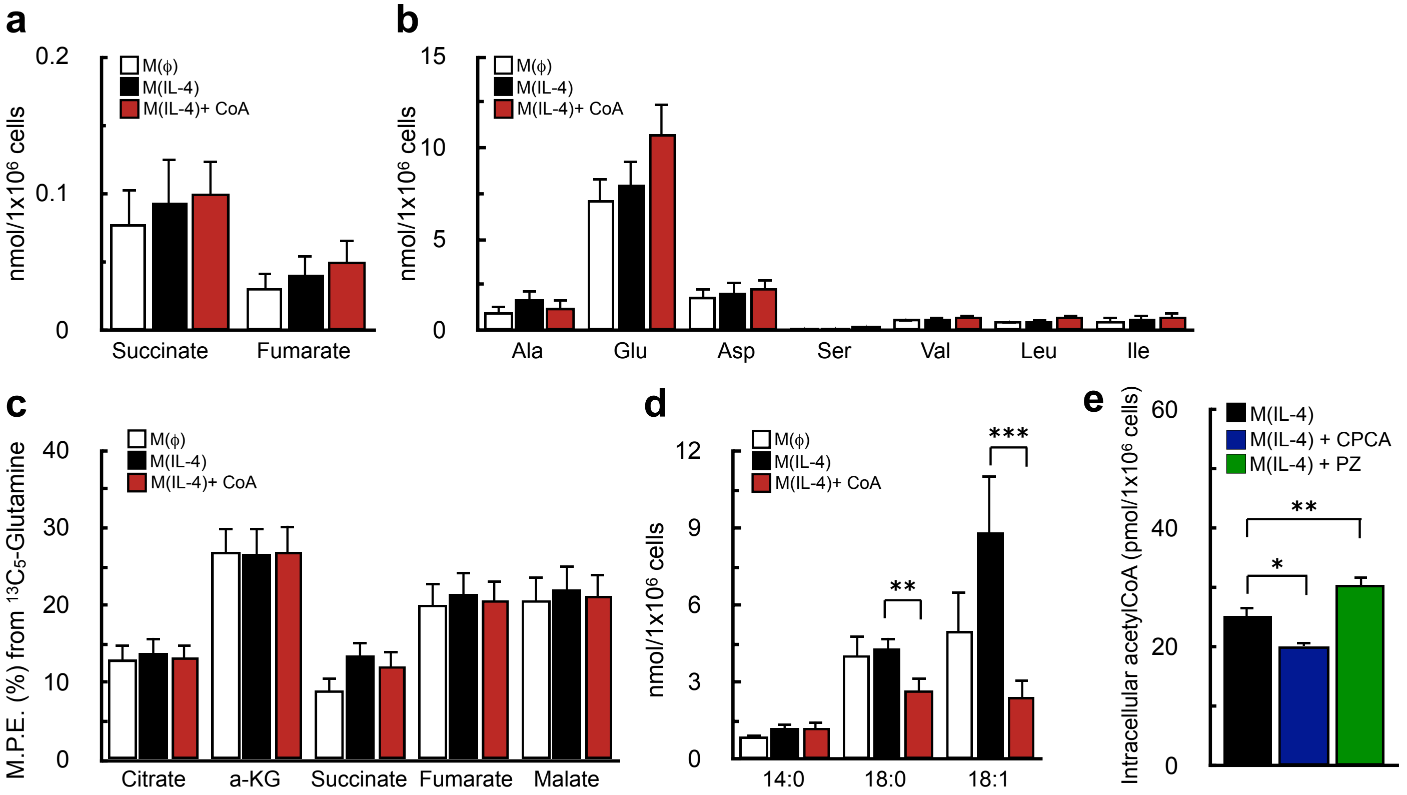


**Supplemental Figure S2. CoA does not enhance alternative activation by increasing metabolic changes associated with the IL-4-response (a-b)** Abundances of (a) TCA cycle intermediates and (b) amino acids in BMDMs treated with vehicle control, IL-4 (20 ng/mL), or IL-4 + 1 mM CoA for 48 hr. (n=7 independent biological replicates). **(c)** Enrichment from uniformly labeled $^{13}C_5$-glutamine into the TCA cycle intermediates citrate, α-ketoglutarate (a-KG), succinate, fumarate, and malate in BMDMs treated as in (a) (n=7 independent biological replicates).**(d)** Quantification of newly synthesized myristic acid (14:0), stearic acid (18:0), and oleic acid (18:1) from BMDMs treated as in (a) (data shown as n=8 technical replicates from n=2 independent biological replicates). **(e)** Intracellular acetyl CoA levels of BMDMs stimulated with IL-4 (20 ng/mL), IL-4+ CPCA (1 mM), or IL-4 +PZ-2891(10 μM) for 48 hr. (n=5 independent biological replicates). All data are presented as mean ± SEM. *$p < 0.05$; **$p < 0.01$; ***$p < 0.001$.

### Supplemental Information Figure S3

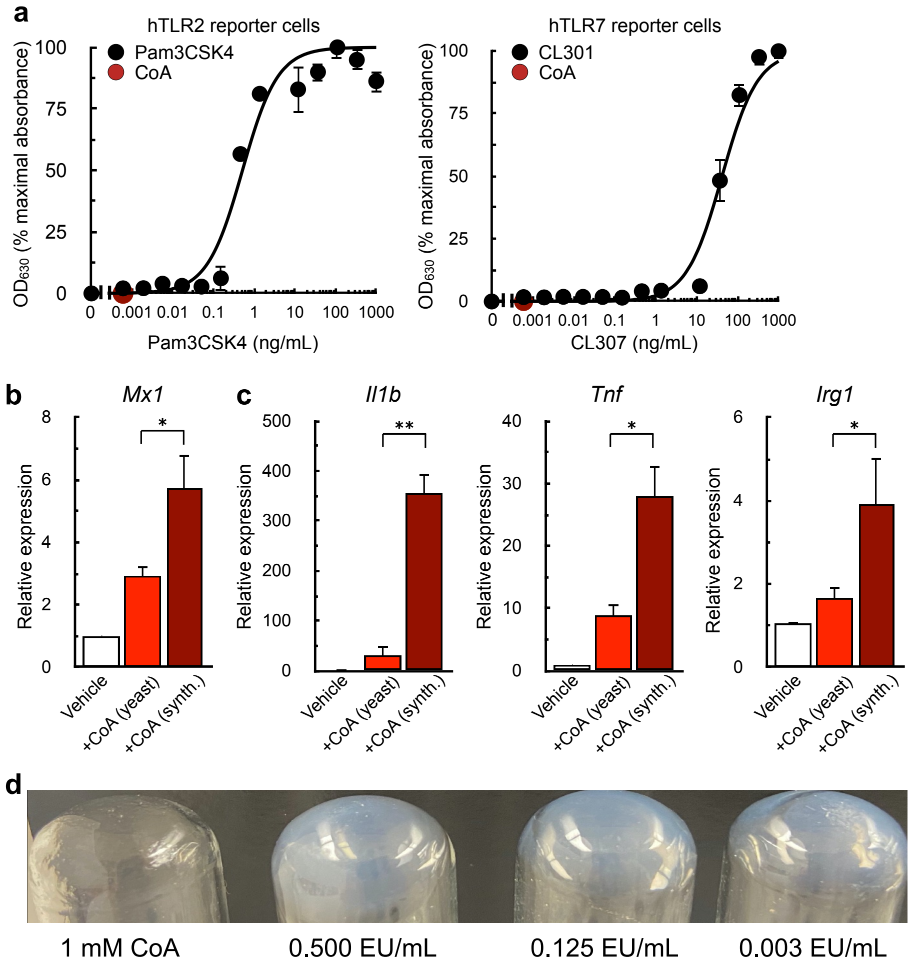


**Supplemental Figure S3. CoA is a TLR4 agonist (a)** Concentration-response curves of hTLR2 and hTLR7 reporter cells exposed to increasing concentrations of Pam3CSK4 and CL301, respectively (black dots). The red dot indicates the response elicited from 1 mM CoA. (n=4 independent biological replicates). **(b)** qPCR analysis of *Mx1* in BMDMs stimulated for 24 hr. with 1 mM CoA that was either chemically synthesized or isolated from yeast (n=5 independent biological replicates). **(c)** qPCR analysis of *Il1b*, *Tnf*, and *Irg1* in BMDMs stimulated for 4 hr. with CoA as in (c) (n=5 independent biological replicates). **(d)** Limulus test, with opaque precipitate signifying the presence of endotoxin, for 1 mM yeast-derived CoA alongside endotoxin standards. The translucent tube shows that CoA is free of endotoxin (EU, endotoxin units). All data are presented as mean ± SEM. $*p < 0.05$; $**p < 0.01$.

**Supplemental Information Figure S4**

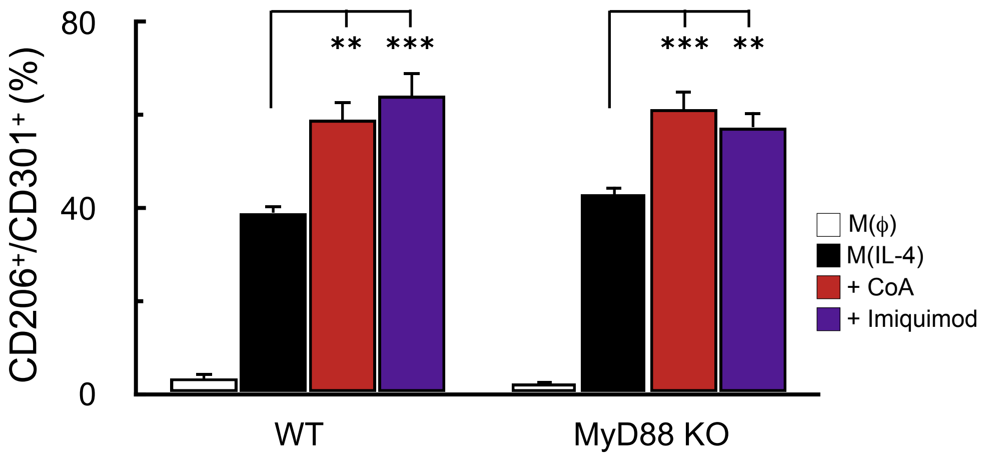


**Supplemental Figure S4. CoA and imiquimod enhance IL-4 cell surface markers despite loss of MyD88**
Percentage of $CD206^+/CD301^+$ in BMDMs harvested from WT and *Myd88*$^{-/-}$ animals upon treatment with vehicle control, IL-4 (20 ng/mL), CoA (1mM), or imiquimod (10μM). (n=7 biological replicates) All data are presented as mean ± SEM. **p < 0.01; ***p<0.001.

**Supplemental Information Figure S5**

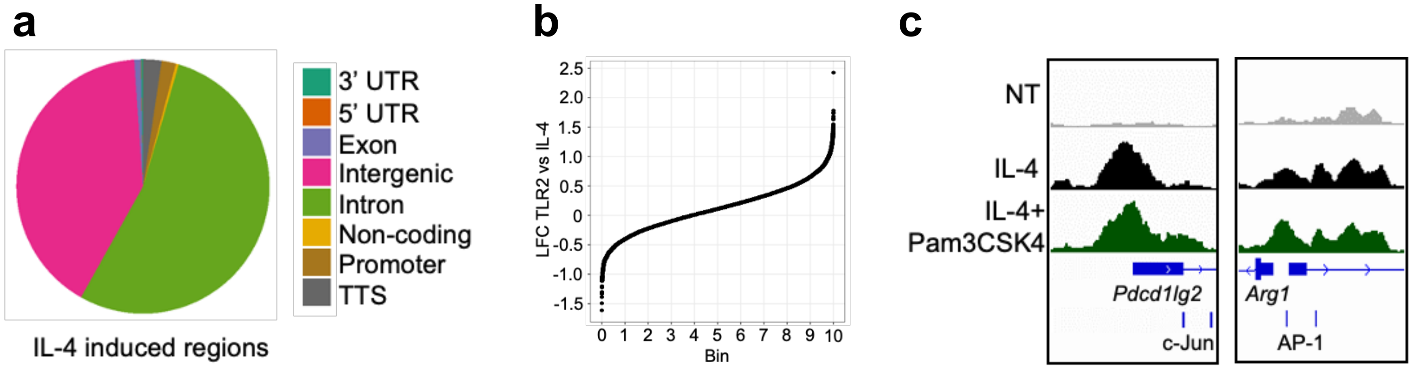


**Supplemental Figure S5. Pam3CSK4 increases the chromatin accessibility of IL-4-induced regions (a)** Genomic distribution of the 10,878 IL-4-induced accessibility locations. **(b)** Distribution of Pam3 and IL-4 co-treatment on the IL-4 inducible regions, divided into 10 equal bins. **(c)** Representative tracks of the alternative activation genes *Pdc1lg2* and *Arg1* promoter regions with nearby c-Jun/AP-1 motifs.
